# Viral mimicry of p65/RelA transactivation domain to inhibit NF-κB activation

**DOI:** 10.1101/2020.10.23.353060

**Authors:** Jonas D. Albarnaz, Hongwei Ren, Alice A. Torres, Evgeniya V. Shmeleva, Carlos A. Melo, Andrew J. Bannister, Matthew P. Brember, Betty Y.-W. Chung, Geoffrey L. Smith

**Affiliations:** Department of Pathology, University of Cambridge, Cambridge CB2 1QP, UK; The Gurdon Institute, University of Cambridge, Cambridge CB2 1QN, UK

**Keywords:** NF-κB, p65, F14, molecular mimicry, CBP, BRD4, poxvirus, vaccinia virus, immune evasion, virulence, HPV16 E7, HSV-1 VP16

## Abstract

Sensing of virus infection activates NF-κB to induce the expression of interferons, cytokines and chemokines to initiate the antiviral response. Viruses antagonise these antiviral defences by interfering with immune sensing and blocking the actions of antiviral and inflammatory molecules. Here, we show that a viral protein mimics the transactivation domain of the p65 subunit of NF-κB. The C terminus of vaccinia virus (VACV) protein F14 (residues 51-73) activates transcription when fused to a DNA-binding domain-containing protein and F14 associates with NF-κB co-activator CBP, disrupting p65-CBP interaction. Consequently, F14 diminishes CBP-mediated acetylation of p65 and the downstream recruitment of the transcriptional regulator BRD4 to the promoter of the NF-κB-responsive genes *CXCL10* and *CCL2*, hence inhibiting their expression. Conversely, the recruitment of BRD4 to the promoters of *NFKBIA*, which encodes the inhibitor of NF-κB (IκBα), and *CXCL8* remains unaffected in the presence of either F14 or JQ1, a competitive inhibitor of BRD4 bromodomains, indicating its recruitment is acetylation-independent. Therefore, unlike other viral NF-κB antagonists, F14 is a selective inhibitor of NF-κB-dependent gene expression. A VACV strain lacking F14 showed that it contributes to virulence in an intradermal model of infection. Our results uncover a mechanism by which viruses disarm the antiviral defences through molecular mimicry of a conserved host protein and provide insight into the regulation of NF-κB-dependent gene expression by BRD4.

## INTRODUCTION

Viruses provide constant selective pressure shaping the evolution of the immune systems of multicellular organisms [1–3]. At the cellular level, an array of receptors detects virus-derived molecules, or more broadly pathogen-associated molecular patterns (PAMPs), allowing the recognition of invading viruses and the activation of a gene expression programme that initiates the antiviral response [reviewed by [4, 5]]. The induced gene products, which include interferons, cytokines and chemokines, are secreted and function as signals to activate more specialised immune cells and attract them to the site of infection, thereby generating inflammation [reviewed by [6–8]]. This coordinated inflammatory response evolved to achieve the control and (or) elimination of the infection, and the establishment of an immunological memory against future infection [reviewed by [4, 9]].

Engagement of pattern recognition receptors (PRRs) by their cognate PAMPs activates multiple transcription factors, including nuclear factor kappa light-chain enhancer of activated B cells (NF-κB) [reviewed by [10–12]]. NF-κB is a homo- or heterodimer of Rel proteins, with the heterodimer of p50 (also known as NF-κB1 or NFKB1) and p65 (also known as RelA or RELA) being the prototypical form of NF-κB [13]. Through an interface formed by the Rel homology domains of the two Rel subunits, NF-κB recognises and binds to a consensus DNA sequence in the promoter elements and enhancers of target genes [reviewed by [10], [14]]. NF-κB-responsive gene products include inflammatory mediators, such as cytokines, chemokines and cell adhesion molecules, as well as proteins involved in other immune processes, like MHC molecules, growth factors and regulators of apoptosis [15, 16]. Moreover, cytokines, such as interleukin (IL)-1 and tumour necrosis factor (TNF)-α, also trigger NF-κB activation upon engagement of their receptors on the cell surface [reviewed by [10, 11]].

In resting conditions, NF-κB remains latent in the cytoplasm bound to the inhibitor of κB (IκB) α (also known as NFKBIA) [17, 18]. Upon activation, the IκB kinase (IKK) complex phosphorylates IκBα, triggering its ubiquitylation and subsequent proteasomal degradation, thus releasing NF-κB to accumulate in the nucleus [18] [reviewed by [10, 11]]. In the nucleus, NF-κB interacts with chromatin remodelling factors, coactivators and general transcription factors to activate the transcription of antiviral and inflammatory genes by RNA polymerase (RNAP) II [14, 19–24]. The specificity and kinetics of NF-κB-dependent gene expression is determined by several factors including dimer composition [25], cooperation with other transcription factors [24], duration of the stimulus [26, 27], cell type [28, 29] and chromatin context on the promoters of target genes [14, 30, 31]. In addition, NF-κB undergoes multiple posttranslational modifications, in the cytoplasm or nucleus, that control its transcriptional activity through interactions with coactivators and basal transcription machinery [reviewed by [13, 32, 33]].

Following the stimulation with PAMPs or inflammatory cytokines (e.g., TNF-α), two conserved residues in p65 are phosphorylated: S276, mainly by protein kinase A (PKA), and S536 within the transactivation domain (TAD), by the IKK complex [34, 35]. Phosphorylation of either site enhances NF-κB transcriptional activity by promoting the interaction with the coactivators CREB-binding protein (CBP) or its paralogue p300 (also known as CREBBP and EP300, respectively). These coactivators acetylate both p65 at K310 and histones on the target gene promoters to allow transcription initiation and elongation to proceed [35–38]. The bromodomain and extraterminal domain (BET) protein BRD4 docks onto acetylated p65-K310, via its two bromodomains, and subsequently recruits positive transcription elongation factor b (P-TEFb) to drive transcription of inflammatory genes by RNAP II [39]. This latter study highlighted the complexity of the gene expression programme downstream of NF-κB, with subsets of genes differentially expressed depending on the transcriptional regulatory events following NF-κB recruitment to DNA [[39–41]; reviewed by [42]]. Targeting of the nuclear activity of NF-κB and its coactivator CBP by viral proteins has been described as a strategy to antagonise antiviral responses (e.g. high-risk human papillomavirus (HPV) E6 protein [43] and herpes simplex virus (HSV) type 1 protein VP16 [44]). However, these studies do not elucidate in detail how viral interference with NF-κB in the nucleus affects induction of the inflammatory genes by this transcription factor. Furthermore, there are contradictory reports regarding the interaction between VP16 and CBP [21, 44].

The confrontations between viruses and hosts leave genetic signatures over their evolutionary histories [3, 45]. On one hand, host innate immune factors display strong signs of positive selection to adapt to the pressure posed by viruses [reviewed by [46]]. On the other hand, viruses acquire multiple mechanisms to antagonise host innate immunity, such as mimicking host factors to disrupt their functions in the antiviral response or to subvert them for immune evasion [reviewed by [46, 47]]. Poxviruses have been a paradigm in the study of virus-host interactions [reviewed by [48]]. Their large DNA genomes encode a plethora of proteins that antagonise the host antiviral response. Some poxvirus proteins show structural similarity to host proteins and modulate innate immune signalling during infection [reviewed by [49, 50]]. For instance, vaccinia virus (VACV), the smallpox vaccine and the prototypical poxvirus, encodes a family of proteins sharing structural similarity to cellular Bcl-2 proteins despite very limited sequence similarity. Viral Bcl-2-like proteins have evolved to perform a wide range of functions, such as inhibition of NF-κB activation [reviewed by [51]]. VACV protein A49, notwithstanding its Bcl-2 fold, also mimics the IκBα phosphodegron that is recognised by the E3 ubiquitin ligase β-TrCP, thereby blocking IκBα ubiquitylation [52, 53]. Upon NF-κB activation, the IKK complex phosphorylates A49 to create the complete phosphodegron mimic that then engages β-TrCP to prevent IκBα ubiquitylation [54].

Despite the existence of multiple inhibitors of NF-κB from VACV, virus strains lacking individual inhibitors have reduced virulence in mouse models, arguing against their functional redundancy [reviewed by [55, 56]]. Therefore, the detailed study of the mechanisms underpinning the antagonism of NF-κB by VACV and other poxviruses offers an opportunity to dissect the signalling pathways leading to NF-κB activation and their relative contributions to antiviral immunity. Previous work from our laboratory predicted that VACV encodes additional inhibitors of NF-κB because a mutant VACV strain (vv811ΔA49) lacking the function of all known inhibitors of NF-κB still suppresses NF-κB-dependent gene expression without preventing NF-κB translocation to the nucleus [57]. Here, we mapped this NF-κB inhibitory activity to the open reading frame (ORF) *F14L*, which is conserved across all orthopoxviruses, including the human pathogens variola, monkeypox and cowpox viruses. We show that the orthopoxvirus protein F14 inhibits NF-κB and a VACV strain lacking F14 has reduced virulence in a mouse model. Mechanistically, the F14 C terminus mimics the TAD of p65 and outcompetes p65 for binding to the coactivator CBP. As a consequence, F14 reduces p65 acetylation and downstream molecular events required for the activation of a subset of NF- κB-responsive inflammatory genes. The selectivity of F14 inhibition makes it unique among known viral antagonists of NF-κB. The dissection of the mechanism of action of F14 also revealed that the transcriptional regulator BRD4 can be recruited to the chromatin in an acetylation-independent manner.

## RESULTS

### Vaccinia virus protein F14 is a virulence factor that inhibits NF-κB activation

The prediction that VACV encodes additional inhibitor(s) of NF-κB that function downstream of p65 translocation to the nucleus [57] prompted a search for nuclear NF-κB inhibitors. VACV strain vv811ΔA49 lacks 56 ORFs, but retains the inhibitor(s) [57, 58], and so its encoded proteins were screened by bioinformatics for ones that met the following criteria: (i) early expression, based on previous VACV transcriptome studies [59, 60]; (ii) predicted not to be involved in replication and/or morphogenesis; (iii) being poorly characterised; (iv) the presence of putative nuclear localisation signals (NLS) or predicted molecular mass <40 kDa that would allow diffusion into the nucleus [61]; and (v) the presence of domains indicative of function. The genomic position of the ORF and its conservation among orthopoxviruses were also considered given that VACV immunomodulatory genes are located towards the genome termini and often are less conserved than genes having functions in virus replication [62].

This approach yielded a list of seven ORFs, namely *F6L*, *F7L*, *F14L*, *A47L*, *B6R*, *B11R* and *B12R*. The proteins encoded by these ORFs were tested for inhibition of NF-κB activation in an NF-κB luciferase reporter gene assay. GFP and VACV protein N2, an interferon regulatory factor (IRF) 3 inhibitor [63], were used as negative controls, whereas VACV protein B14, a known inhibitor of NF-κB [64], was used as positive control. Protein F14 inhibited TNF-α- and IL-1β-stimulated NF-κB activity in HEK 293T cells in a dose-dependent manner (Figure 1A, B and Figure S1). This inhibitory activity was specific for the NF-κB pathway, because F14 did not affect IFN-α-stimulated IFN-α/β receptor (IFNAR)/signal transducer and activator of transcription (STAT) or phorbol 12-myristate 13-acetate (PMA)-stimulated mitogen-activated protein kinase (MAPK)/AP-1 pathways (Figure 1C, D). The inhibitory activity was exerted despite the lower levels of F14 when compared to protein B14 (Figure 1E) [64]. Conversely, VACV protein C6 suppressed type I IFN signalling and B14 upregulated AP-1 activity, as observed previously (Figure 1C, D) [65, 66].

**Figure 1:**
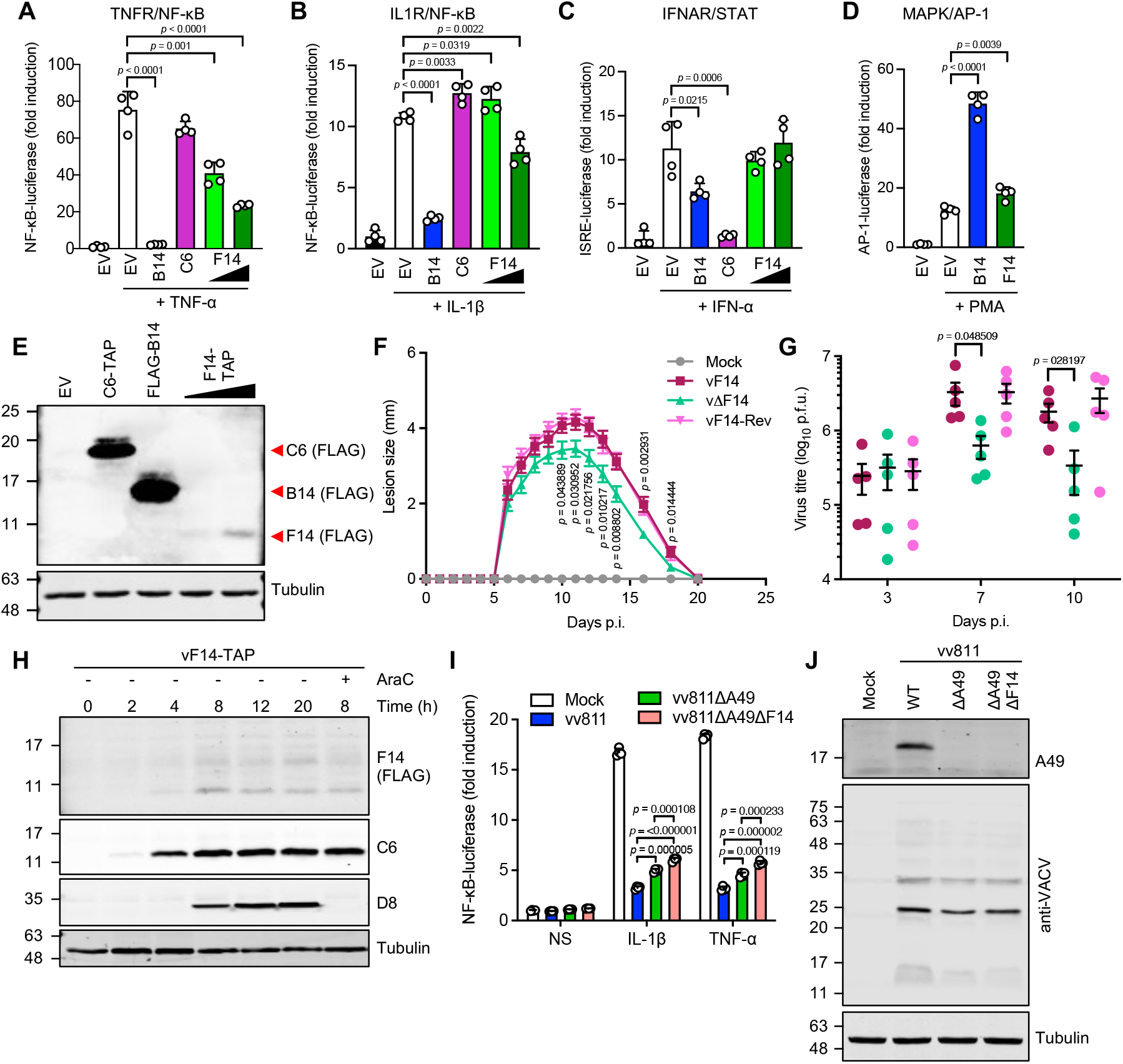
Vaccinia virus protein F14 inhibits NF-κB activation and contributes to virulence. (**A-D**) NF-κB- (**A, B**), IFNAR/STAT- (**C**), or MAPK/AP-1- (**D**) dependent luciferase activities in HEK 293T cells transfected with vectors expressing the indicated VACV proteins or empty vector (EV), and stimulated with TNF-α, IL-1β, IFN-α or PMA (as indicated). Means + s.d. (*n* = 4 per condition) are shown. (**E**) Immunoblotting of whole cell lysates of transfected HEK 293T cells. (**F**) C57BL/6 mice were infected intradermally in both ears with 10^4^ p.f.u. of the indicated VACV strains and the lesions were measured daily. Means ± s.e.m. (*n* = 10 mice) are shown. (**G**) Virus titres in the ears of mice infected as in (**F**). Means ± s.e.m. (*n* = 5 mice) are shown. (**H**) Immunoblotting of protein extracts from HeLa cells infected with vF14-TAP (5 p.f.u./cell) and treated with cytosine arabinoside (AraC, 40 µg/ml) where indicated. (**I**) NF-κB- dependent luciferase activity in A549 cells infected with VACV vv811 strains (5 p.f.u./cell, 6 h) and stimulated with TNF-α or IL-1β for additional 6 h. Means + s.d. (*n* = 4 per condition) are shown. (**J**) Immunoblotting of protein extracts from A549 cells infected as in **(I)**. The positions of molecular mass markers in kDa are shown on the left of immunoblots. Immunoblots of tagged proteins are labelled with the protein name followed the epitope tag antiboby in parentheses. When multiple tagged proteins are shown in the same immunoblot, each protein is indicated by a red arrowhead. Statistical significance was determined by the Student’s *t*-test. Data shown in (**A-D, I**), (**F, G**) and (**E, H, J**) are representative of four, two or three separate experiments, respectively.

The virulence of VACV strains lacking specific genes has been tested mostly in intranasal or intradermal murine models [reviewed by [55, 56]]. Deletion of genes encoding VACV immunomodulatory proteins may give a phenotype in either, neither or both models. To evaluate if loss of F14 expression affected virulence, a recombinant VACV lacking F14 was generated, termed vΔF14. Intradermal injection of the ear pinnae of mice with vΔF14 produced smaller lesions, and reduced virus titres at 7 and 10 d post-infection (p.i.) (Figure 1F, G) compared to wildtype virus (vF14) and a revertant strain (vF14-Rev), generated by reinserting F14 into vΔF14 at its natural locus (Figure 1F, G). Attenuation of vΔF14 in the intradermal mouse model correlated with reduced viral titres in the infected ears 7 and 10 d p.i., but not 3 d p.i. (Figure 1G). In contrast, in an intranasal mouse model, vΔF14 caused the same extent of body mass loss as wildtype and revertant controls (Figure S2). In cell culture, vF14, vΔF14 and vF14-Rev displayed no differences in replication and plaque size (Figure S3). Altogether, these experiments showed that F14 is not essential for virus replication but contributes to virulence.

Previous analyses of the VACV transcriptome showed that the *F14L* ORF is transcribed and translated early during infection [59, 60, 67]. This was consistent with an upstream typical early promoter and a transcription termination motif T5NT downstream of the stop codon [68, 69]. To investigate F14 expression during infection, a VACV strain was constructed in which F14 was tagged with a C-terminal TAP tag. The vF14-TAP strain replicated normally (Figure S3) and immunoblotting showed F14 protein expression was detected from 4 h p.i. and peaked by 8 h p.i., matching the accumulation of the early VACV protein C6 (Figure 1H) [70]. F14 levels were notably low either when expressed ectopically (Figure 1E) or during infection (Figure 1H). This might explain why F14 was not detected in our recent quantitative proteomic analysis of VACV infection, which detected about 80% of the predicted VACV proteins [71]. Pharmacological inhibition of virus DNA replication with cytosine arabinoside (AraC) did not affect F14 protein levels, consistent with early expression, whereas late protein D8 was inhibited (Figure 1H).

The existence of multiple VACV inhibitors of NF-κB that each contribute to virulence indicates they are not redundant. To test if F14 affects NF-κB activation during infection, we deleted F14 from the vv811ΔA49 strain that lacks other known inhibitors of NF-κB [57] and then infected an NF-κB reporter cell line [57]. As shown previously, vv811ΔA49 inhibited NF-κB to a reduced extent when compared to the parental vv811 strain (Figure 1I) [57] and deletion of *F14L* from vv811ΔA49 reduced NF-κB inhibition further (Figure 1I). Immunoblotting confirmed equal infection with these viruses (Figure 1J). Notably, vv811ΔA49ΔF14 still suppressed NF- κB activation considerably, which might be explained by: (i) the existence of additional virally-encoded inhibitors that cooperate to inhibit NF-κB in the nucleus, or (ii) the actions of D9 and D10 decapping enzymes to reduce host mRNA [72, 73].

### F14 inhibits NF-κB at or downstream of p65

To dissect how F14 functions, its impact on three hallmarks of NF-κB signalling were studied: namely, degradation of IκBα, phosphorylation of p65 at S536 and p65 nuclear translocation. A cell line that expresses F14 inducibly upon addition of doxycycline was used to study the degradation of IκBα, and phosphorylation and nuclear translocation of p65 following stimulation with TNF-α. IκBα degradation was evident 15 min after stimulation and its re- synthesis had started by 30 min, but neither process was influenced by F14 (Figure 2A). F14 also did not affect phosphorylation of p65 at S536 (Figure 2A) or p65 translocation into the nucleus as measured by immunofluorescence (Figure 2B and Figure S4). In contrast, VACV protein B14 inhibited translocation efficiently as reported [64], and VACV protein C6, an IFN antagonist [65, 70], did not (Figure 2B).

**Figure 2:**
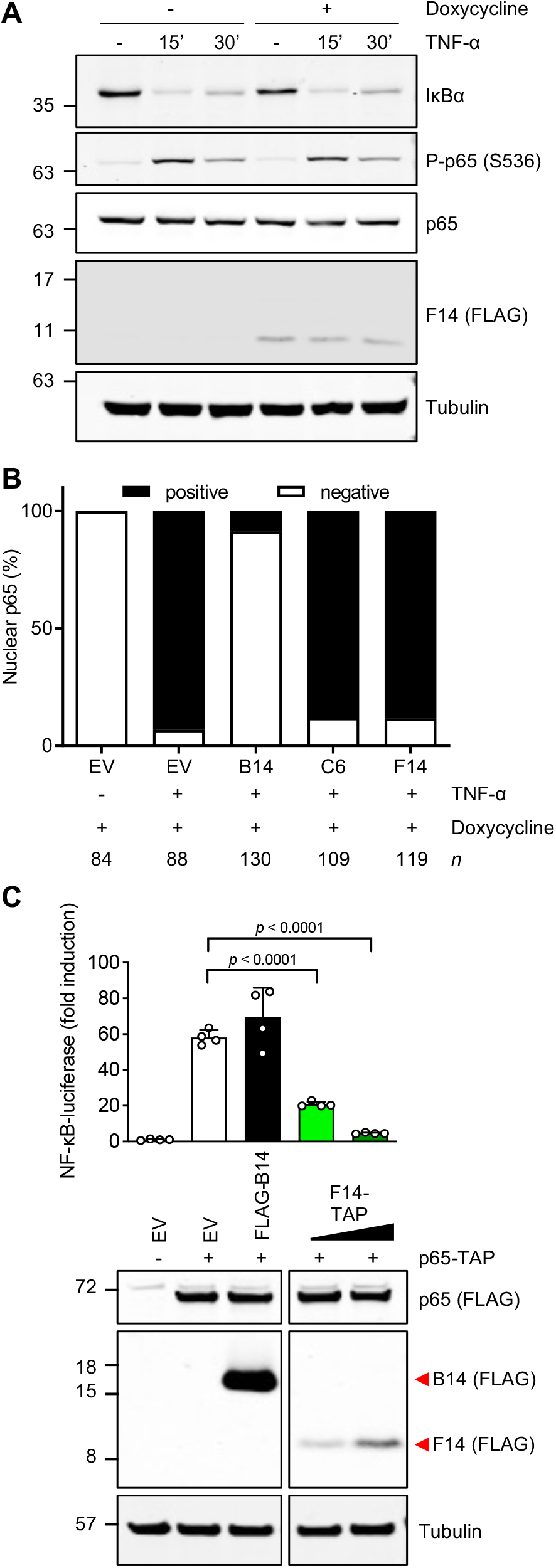
F14 inhibits NF-κB at or downstream of p65. (**A**) Immunoblotting of protein extracts from T-REx-293 cells inducibly expressing F14, after doxycycline induction overnight and TNF-α stimulation. Data are representative of three independent experiments. (**B**) Quantification of NF-κB p65 localisation after immunofluorescence of T-REx-293 cells inducibly expressing the empty vector (EV) or VACV proteins B14, C6 or F14, induced with doxycycline and stimulated with TNF-α for 15 min. Number of cells counted from two independent experiments (*n*) is stated below each bar. (**C**) NF-κB activity in HEK 293T cells transfected with vectors expressing p65, VACV proteins B14 or F14, or empty vector (EV). Top panel: Means + s.d. (*n* = 4 per condition) are shown. Statistical significance was determined by Student’s *t*-test. Bottom panel: Immunoblotting. Protein molecular mass markers in kDa are shown on the left of the blots. Immunoblots of tagged proteins are labelled with the protein name followed by the epitope tag antiboby in parentheses. When multiple tagged proteins are shown in the same immunoblot, each protein is indicated by a red arrowhead.

Next, the NF-κB inhibitory activity of F14 was tested by reporter gene assay following pathway activation by p65 overexpression. In contrast to B14, F14 inhibited p65-mediated activation in a dose-dependent manner without affecting p65 levels (Figure 2C). Altogether, these results showed that F14 blocks NF-κB in the nucleus at or downstream of p65. F14 thus fits the criteria described previously for the unknown inhibitor of NF-κB encoded by VACV and expressed by vv811ΔA49 [57].

### F14 orthologues are conserved in orthopoxviruses and mimic the p65 transactivation domain

Poxvirus immunomodulatory proteins are generally encoded in the variable genome termini, share lower sequence identity and show genus-specific distribution [62]. Even among orthopoxviruses, only a few of the immunomodulatory genes are present in all virus species [62]. Nonetheless, Viral Orthologous Clusters [74] and protein BLAST searches found orthologues of VACV F14 in all orthopoxviruses, with 70.7% to 98.6% amino acid (aa) identity (Figure 3A). The *F14L* orthologue from cowpox virus, *CPXV062*, is expressed during infection by two different strains of cowpox virus, as shown by RNA sequencing [75]. *F14L* orthologues from monkeypox and variola viruses are also likely to be expressed during infection, because the nucleotide sequences surrounding the ORFs contained highly conserved transcriptional regulatory sequences, i.e., canonical poxvirus early promoters and transcription termination signals [68]. Furthermore, historical strains of variola virus from the 10^th^ and 17^th^ centuries CE also encoded *F14L* orthologues with conserved flanking transcription regulatory sequences [76, 77].

**Figure 3:**
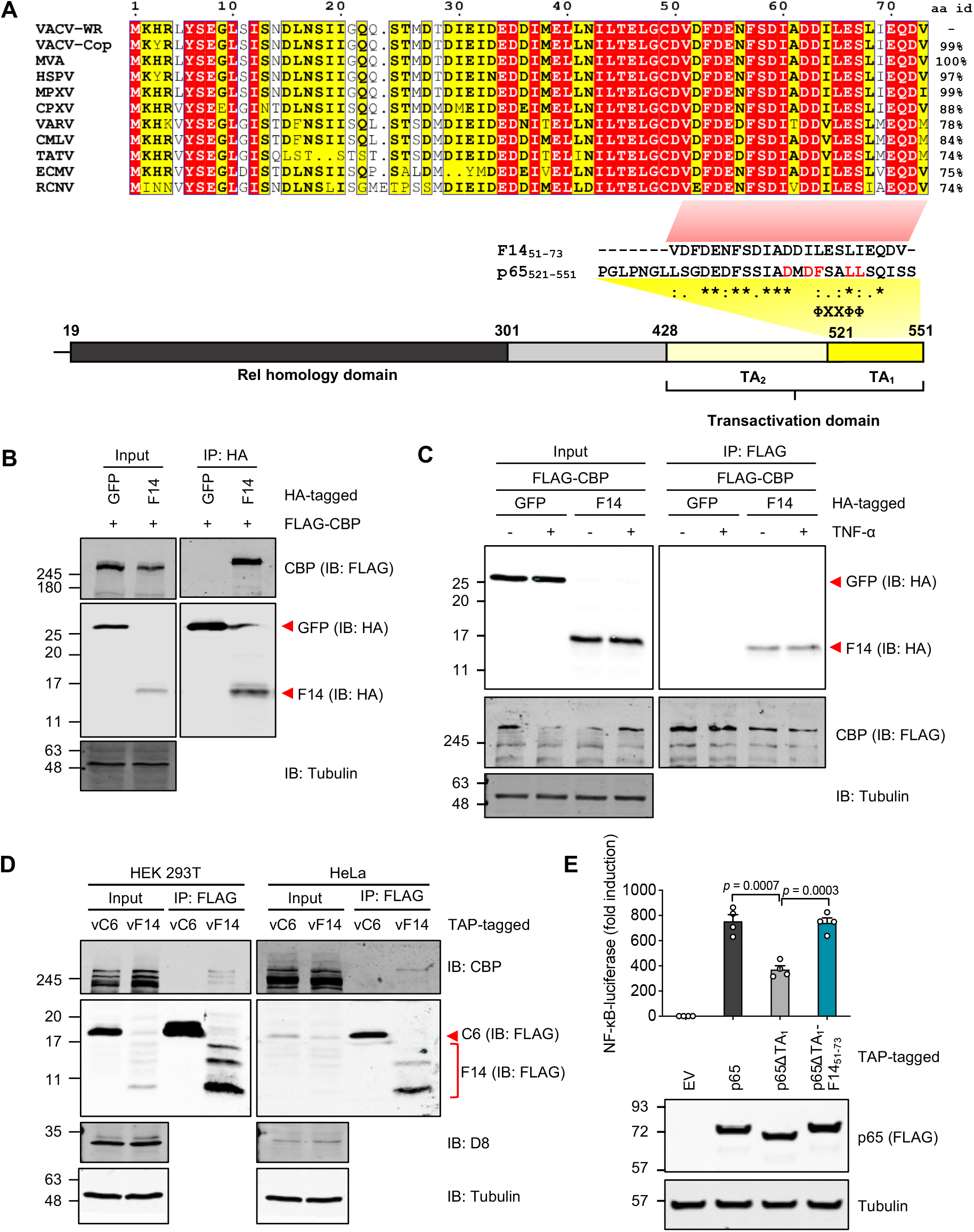
F14 binds to CBP and has transactivation activity. (**A**) Top: Amino acid alignment of F14 orthologues of representative orthopoxviruses: vaccinia virus (VACV) Western Reserve (WR), VACV Copenhagen (Cop), modified vaccinia Ankara (MVA), horsepox virus (HSPV), monkeypox virus (MPXV), cowpox virus (CPXV), variola virus (VARV), camelpox virus (CMLV), taterapox virus (TATV), ectromelia virus (ECMV), and racoonpox virus (RCNV). Red, aa identical in all sequences; yellow, aa identical in at least 8/11 sequences. The percent aa identity of F14 orthologues compared to the F14 protein of VACV-WR are shown on the right. Bottom: Alignment of the C-termini of F14 and p65 highlighting their sequence similarity including the ϕXXϕϕ motif, above a schematic of p65 and its functional domains. Asterisks (*), identical aa; colons (:), conservative aa change; dots (.), non-conservative aa change. Nucleotide sequences used for this study are listed in Table S1. (**B, C**) Lysates from transfected HEK 293T cells were immunoprecipitated with anti-HA (**B**) or anti-FLAG (**C**). Immunoblots are representative of three independent experiments. (**D**) HEK 293T and HeLa cells were infected with VACV strains vC6-TAP or vF14-TAP (5 p.f.u./cell, 8 h) and lysates were immunoprecipitated with anti-FLAG. Immunoblots are representative of two independent experiments. (**E**) NF-κB-dependent luciferase activity in HEK 293T cells transfected with vectors expressing p65, p65 mutants or empty vector (EV). Top panel: Means + s.d. (*n* = 4 per condition) are shown. Statistical significance was determined by the Student’s *t*-test. Bottom panel: Immunoblotting. Protein molecular mass markers in kDa are shown on the left of the blots. Immunoblots of tagged proteins are labelled with the protein name followed the epitope tag antiboby in parentheses. When multiple tagged proteins are shown in the same immunoblot, each protein is indicated by a red arrowhead.

The C-terminal half of F14 was more conserved and included a predicted coiled-coil region (aa 34-47), the only structural motif predicted via bioinformatic analyses. However, the Phyre2 algorithm [78] predicted the C-terminal aa 55 to 71 to adopt an α-helical secondary structure similar to that of aa 534-546 of p65 in complex with the PH domain of human general transcription factor Tfb1, or aa 534-546 of p65 in complex with the KIX domain of NF-κB coactivator CBP [79]. The F14 aa similarity was striking despite the low confidence of the Phyre2 model (41.4%). When aligned to p65 C-terminal aa 521-551, F14 shared 39% aa identity and 61% aa similarity, including the conservation of a ΦXXΦΦ motif and an upstream acidic residue, both essential for NF-κB transcriptional activity [79–81]. The C terminus of p65 harbours its transactivation domain (TAD), which is divided into two subdomains that have independent transcriptional activity: TA1 (aa 521 to 551) and TA2 (aa 428 to 521) [22, 82]. TA1 contributes at least 85% of p65 transcriptional activity and interacts directly with CBP [22, 79, 82]. Notably, in F14 the position equivalent to S536 in p65, which is phosphorylated upon NF- κB activation [34, 38], is occupied by the negatively charged residue D59 (Figure 3A). The negative charge of F14-D59 closely resembles the negative charge conferred by phosphorylation of p65-S536 during NF-κB activation [34].

These observations and the key role of CBP in NF-κB-dependent gene activation [20] prompted investigation of whether F14 could interact with CBP. Immunoprecipitation (IP) of HA-tagged F14 co-precipitated CBP-FLAG from HEK 293T cells (Figure 3B). Reciprocal IP experiments showed that ectopic CBP co-precipitated F14-HA, but not GFP-HA, with or without prior TNF-α stimulation (Figure 3C). These interactions were also seen at endogenous levels in both HEK 293T and HeLa cells infected with vF14-TAP. F14, but not C6, co-precipitated endogenous CBP (Figure 3D).

To test whether the C terminus of F14 mediated transactivation via its binding to CBP, F14 aa 51 to 73 were fused to the C terminus of a p65 mutant lacking the TA1 subdomain of the TAD (ΔTA1) and the fusion protein was tested in a NF-κB reporter gene assay. Compared to wildtype p65, the p65ΔTA1 mutant was impaired in its transactivating activity, which was restored to wildtype levels upon fusion to F1451-73 (Figure 3E). This result argues strongly that the C terminus of F14 mimics the TA1 of p65 and this mimicry might explain how F14 inhibits NF-κB activation.

### F14 outcompetes NF-κB for binding to CBP

The similarity between the C termini of F14 and p65 led us to investigate if conserved aa residues contributed to the NF-κB inhibitory activity of F14. Based on the structure of CBP KIX domain in complex with p65 TA1 [79], residues of the F14 TAD-like domain corresponding to residues of p65 important for its transcriptional activity and binding to CBP were mutated. Three sites were altered by site-directed mutagenesis: the dipeptide D62/63, and the following L65 and L68 of the ΦXXΦΦ motif. F14 L65A or L68A still inhibited NF-κB (Figure 4A), although the L65A mutant was slightly impaired. In contrast, mutation of D62/63 to either alanine (D62/63A) or lysine (D62/63K) abolished the inhibitory activity (Figure 4A). Protein levels were comparable across the different F14 mutants (Figure 4A). The loss of NF-κB inhibitory activity of D62/63A and D62/63K mutants correlated with their reduced capacity to co-precipitate CBP, whereas L65A and L68A mutants co-precipitated CBP to the same extent as wildtype F14 (Figure 4B). The mutation of the negatively charged D62/63 to positively charged lysine residues was more efficient in disrupting the interaction between F14 and CBP than only abolishing the charge (Figure 4B). Collectively, these results highlight the importance of the negatively charged dipeptide D62/63 within the TAD-like domain for NF-κB inhibition by F14.

**Figure 4:**
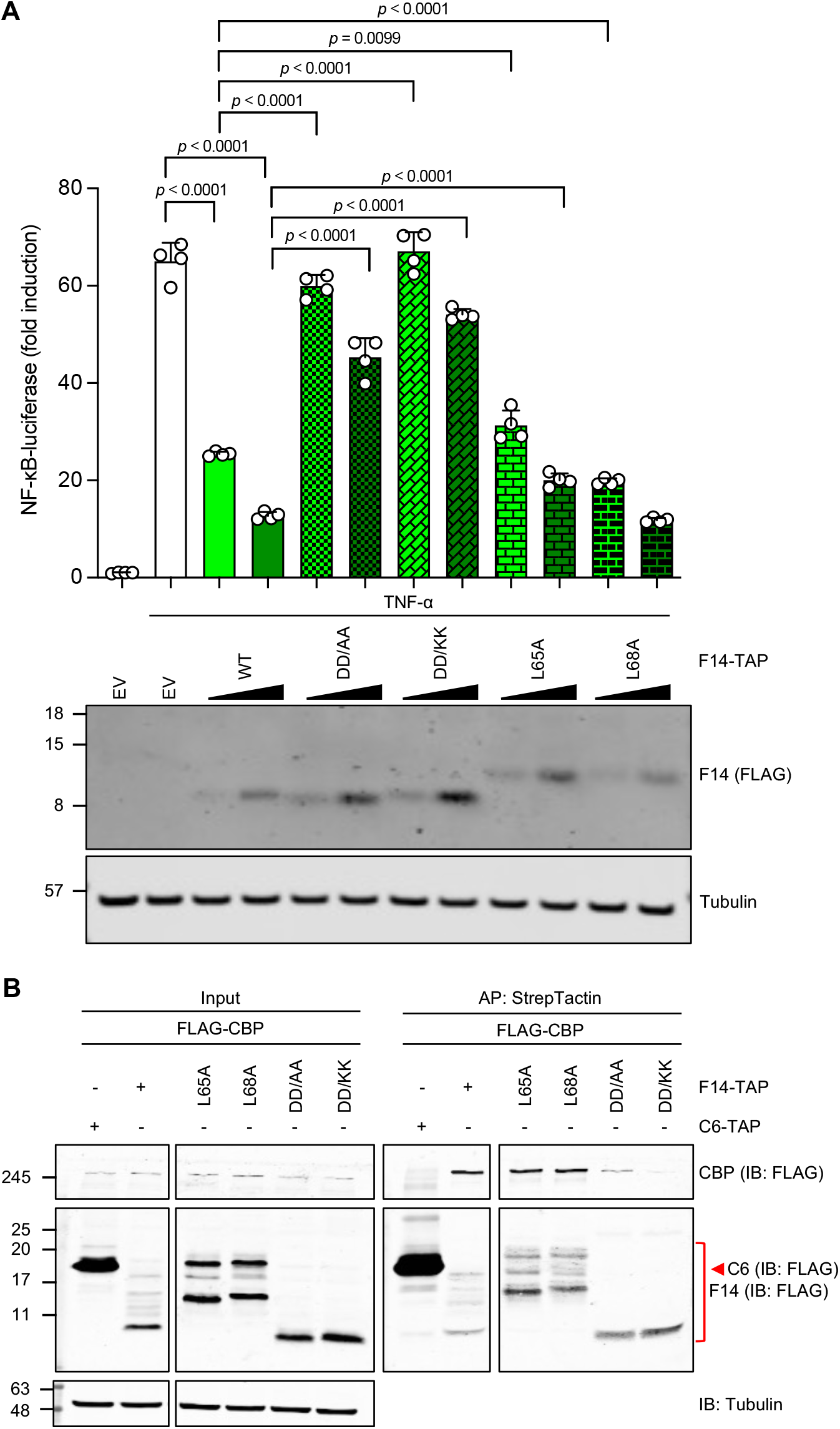
The dipeptide D62/63 of F14 is required for inhibition of NF-κB. (**A**) NF-κB- dependent luciferase activity in HEK 293T cells transfected with vectors expressing F14, F14 mutants, or empty vector (EV), and stimulated with TNF-α for 8 h. Top panel: Means + s.d. (*n* = 4 per condition) are shown. Statistical significance was determined by the Student’s *t*-test. Bottom panel: Immunoblotting. (**B**) Lysates from transfected HEK 293T cells were affinity- purified with StrepTactin resin. DD/AA denotes D62/63A mutant and DD/KK, D62/63K mutant. Protein molecular mass markers in kDa are shown on the left of the blots. Immunoblots of tagged proteins are labelled with the protein name followed the epitope tag antiboby in parentheses. When multiple tagged proteins are shown in the same immunoblot, each protein is indicated by a red arrowhead. Data are representative of three independent experiments.

Next, we tested if F14 could disrupt the interaction of p65 with its coactivator CBP [20]. HEK 293T cells were transfected with vectors expressing p65 and CBP or RIG-I (negative control), and VACV proteins F14 or C6. The amount of p65-HA immunoprecipitated by ectopic CBP was reduced by increasing amounts of F14 but not C6 (Figure 5A, B). Quantitative analysis showed equivalent ectopic CBP immunoprecipitation with or without F14 (Figure 5C). Furthermore, the mutation D62/63K diminished the capacity of F14 to disrupt the interaction of CBP and p65 (Figure 5D). This observation correlated well with the reduced capacity of the D62/63K mutant to co-precipitate CBP (Figure 4B).

**Figure 5:**
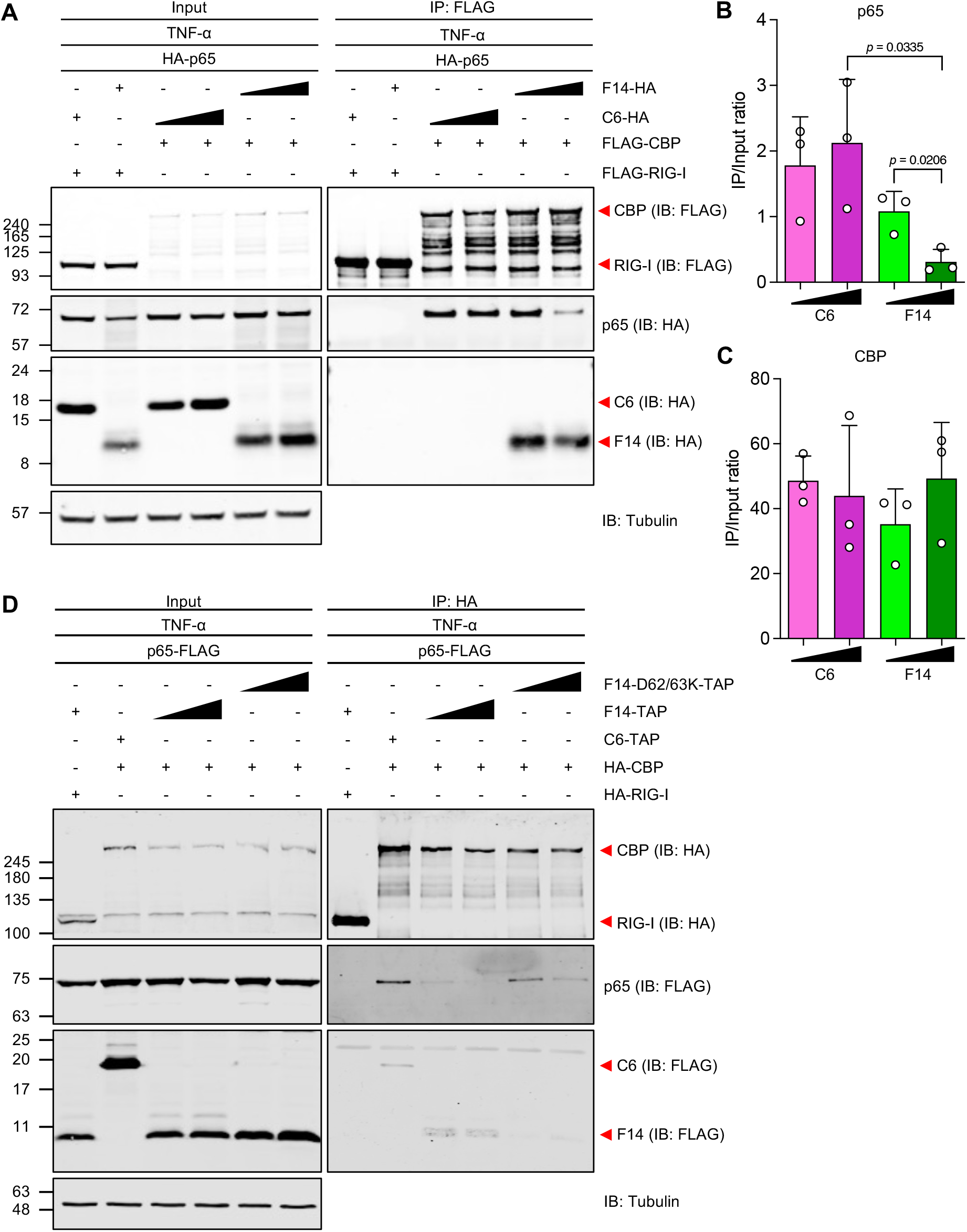
F14 outcompetes NF-κB for binding to CBP. (**A, D**) Lysates from transfected HEK 293T cells were immunoprecipitated with anti-FLAG (**A**) or anti-HA (**D**) after TNF-α stimulation. Immunoblots are representative of three independent experiments. (**B, C**) Ratio of immunoprecipitate (IP) over input signal intensities from immunoblots as in (**A**). Means + s.d. (*n* = 3 independent experiments) are shown. Statistical significance was determined by the Student’s *t*-test. Protein molecular mass markers in kDa are shown on the left of the blots. Immunoblots of tagged proteins are labelled with the protein name followed the epitope tag antiboby in parentheses. When multiple tagged proteins are shown in the same immunoblot, each protein is indicated by a red arrowhead.

### F14 suppresses expression of a subset of NF-κB-responsive genes

To address the impact of F14 on the induction of endogenous NF-κB-responsive genes by TNF-α, the cell line inducibly expressing F14 was utilised. NF-κB-responsive genes display different temporal kinetics upon activation, with “early” gene transcripts peaking between 30 – 60 min after stimulation before declining, whilst “late” gene transcripts accumulate slowly and progressively, peaking 3 h post stimulation [15, 16]. When F14 expression was induced, mRNAs of *NFKBIA* and *CXCL8* “early” genes had equivalent induction kinetics compared to uninduced cells (Figure 6A, B, I). The lack of inhibition of F14 on the expression of *NFKBIA* mRNA is in agreement with the previous finding that the re-synthesis of IκBα (*NFKBIA* protein product) is unaffected by F14 after its proteasomal degradation induced by TNF-α (Figure 2A). Conversely, F14 induction inhibited the accumulation of the mRNAs of *CCL2* and *CXCL10* “late” genes in response to TNF-α (Figure 6D, E). Similar results were observed when the F14-expressing cell line was compared to the cell line inducibly expressing C6 (Figure S5).

**Figure 6:**
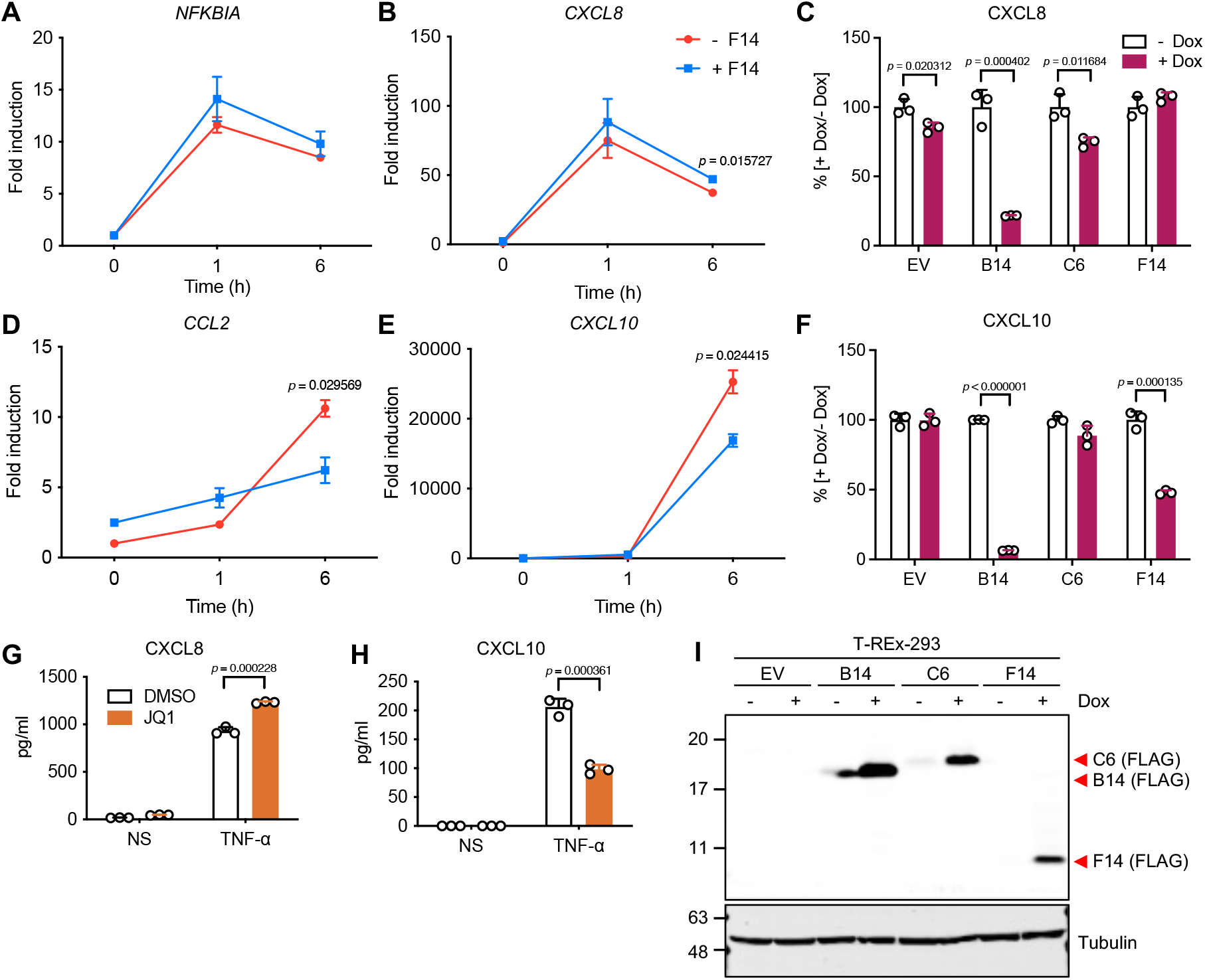
F14 suppresses expression of a subset of NF-κB-responsive genes. **(A, B, D, E)** RT-qPCR analysis of NF-κB-responsive gene expression in inducible T-REx-293-F14 cells in the absence (– F14) or in the presence (+ F14) of doxycycline overnight, and stimulated with TNF-α. Means ± s.d. (*n* = 2 per condition) are shown. (**C, F**) ELISA of culture supernatants from T-REx-293 cells inducibly expressing the empty vector (EV) or VACV proteins B14, C6, or F14, induced overnight with doxycycline and stimulated with TNF-α for 16 h. Means + s.d. (*n* = 3 per condition) of the percent of secretion in presence of doxycycline (+ Dox) versus in the absence of doxycycline (– Dox, equals 100%) are shown. (**G, H**) ELISA of culture supernatants from T-REx-293-EV cells stimulated with TNF-α for 16 h in the absence or in the presence of JQ1. Means ± s.d. (*n* = 3 per condition) are shown. (**I**) Immunoblotting of lysates of inducible T-REx-293 cell lines in the absence or in the presence of doxycycline overnight. Protein molecular masses in kDa are shown on the left of the blots. Immunoblots of tagged proteins are labelled with the protein name followed the epitope tag antiboby in parentheses. When multiple tagged proteins are shown in the same immunoblot, each protein is indicated by a red arrowhead. Statistical significance was determined by the Student’s *t*-test.

*CXCL8* and *CXCL10* encode chemokines CXCL8 and CXCL10 (also known as IL-8 and IP- 10, respectively). Following induction of VACV protein expression, the levels of these secreted chemokines were measured by ELISA and showed that levels of CXCL10, but not CXCL8, was inhibited by F14. In contrast, the secretion of both chemokines was inhibited, or unaffected, by VACV proteins B14 or C6, respectively, as expected (Figure 6C, F, J, I and Figure S6). Thus, unlike other VACV NF-κB inhibitors, F14 is selective and inhibits only a subset of NF-κB-responsive genes.

Differential regulation of transcription activation downstream of NF-κB has been ascribed to the recruitment of BRD4 to some NF-κB-dependent inflammatory genes [39]. Via its bromodomains 1 and 2, BRD4 docks onto acetylated histones and non-histone proteins and recruits transcriptional regulatory complexes to chromatin [reviewed by [83, 84]]. The specific recognition of acetyl-lysine residues by BRD4 is competitively inhibited by small-molecule BET bromodomain inhibitors, such as JQ1 [85]. Therefore, to gain more insight into the mechanism underpinning the selective inhibition of inflammatory genes by F14, the effect of JQ1 on the inducible expression of CXCL8 and CXCL10 was investigated. Following TNF-α stimulation, JQ1 inhibited the secretion of CXCL10, but not CXCL8, phenocopying the selective inhibition of inflammatory protein expression by F14 (Figure 6G, H).

### Acetylation of p65 and recruitment of BRD4 are inhibited by F14

Posttranslational modifications of p65 accompany NF-κB translocation to the nucleus and some, such as acetylation by acetyltransferases CBP and p300, are associated with increased transcriptional activity [[35, 36, 38]; reviewed by [13, 32, 33]]. F14 did not interfere with the phosphorylation of p65 at S536 (Figure 2A), so the acetylation of p65-K310 was investigated. Cell lines that express F14 inducibly or contain the empty vector (EV) control were transfected with plasmids expressing p65 and CBP in the presence of the inducer, doxycycline. Although both cell lines expressed equivalent amounts of ectopic p65 and CBP, the amount of p65 acetylated at K310 was greatly diminished by F14 (Figure 7A). Quantitative analysis showed acetylated p65 was reduced 90% by F14 (Figure 7B). This result, together with data in Figure 5, indicated that the reduced acetylation of p65 was due to disruption of the interaction between p65 and CBP by F14.

**Figure 7:**
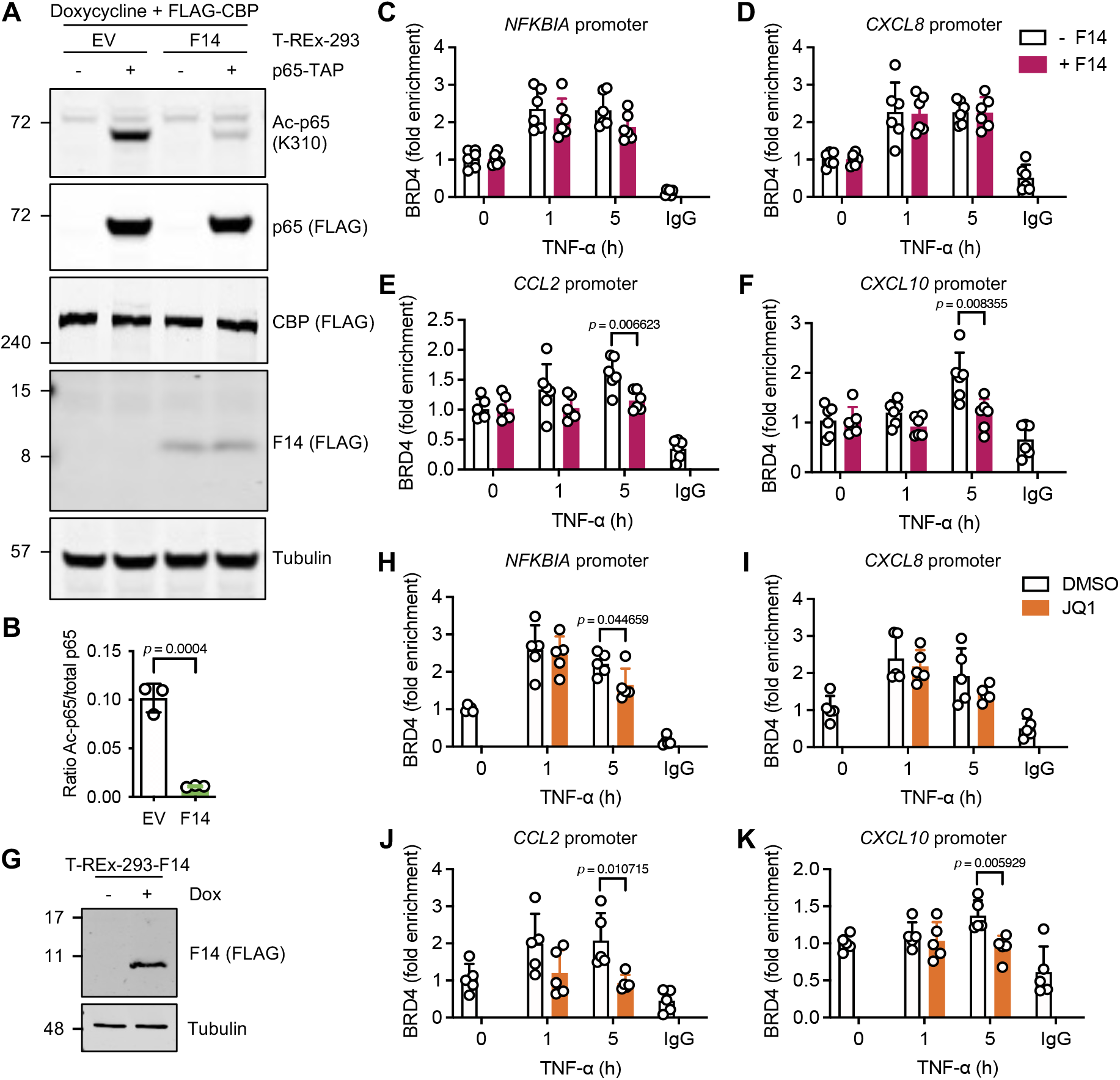
F14 antagonises p65 acetylation and inducible recruitment of BRD4 to *CCL2* and *CXCL10* promoters. (**A**) Immunoblotting of protein lysates from T-REx-293 cells stably transfected with empty vector (EV) or inducibly expressing F14, induced with doxycycline and transfected with plasmids expressing p65 and CBP. Blots are representative of two independent experiments carried out with three biological replicates each. (**B**) Ratio of acetylated (Ac) p65 over total ectopic p65 signal intensities from immunoblots as in (**A**). Means + s.d. (*n* = 3 per condition) are shown. (**C-F, H-K**) Chromatin immunoprecipitation (ChIP) with anti-BRD4 antibody or control IgG, and qPCR for the promoters of *NFKBIA* (**C, H**), *CXCL8* (**D, I**), *CCL2* (**E, J**) and *CXCL10* (**F, K**) genes. T-REx-293-F14 were left uninduced (– F14) or induced with doxycycline (+ F14) and stimulated with TNF-α (**C-F**). Alternatively, T-REx-293 cells were treated with JQ1 before TNF-α stimulation (**H, K**). Means + s.d. (*n* = 5-6 per condition from two independent experiments) are shown. (**G**) Immunoblotting from **(C-F)**. Protein molecular mass markers in kDa are shown on the left of the blots. Immunoblots of tagged proteins are labelled with the protein name followed the epitope tag antiboby in parentheses. When multiple tagged proteins are shown in the same immunoblot, each protein is indicated a red arrowhead. Statistical significance was determined by the Student’s *t*-test.

Acetylated K310 on p65 serves as a docking site for the bromodomains 1 and 2 of BRD4, which then recruits P-TEFb to promote RNAP II elongation during transcription of some NF- κB-responsive genes [39]. The differential sensitivity of TNF-α-stimulated genes to the inhibition of NF-κB by F14 might reflect the differential requirement of p65 acetylated at K310, and the subsequent recruitment of BRD4, to activate the expression from NF-κB-responsive promoters [39]. This hypothesis was tested by chromatin immunoprecipitation with an anti- BRD4 antibody followed by quantitative PCR of the promoters of four representative genes: *NFKBIA* and *CXCL8*, resistant to F14 inhibition, and *CCL2* and *CXCL10*, sensitive to inhibition. BRD4 was recruited to these promoters after TNF-α stimulation, with BRD4 present on *NFKBIA* and *CXCL8* promoters at 1 and 5 h post-stimulation, whereas BRD4 was more enriched on *CCL2* and *CXCL10* promoters only at 5 h post-stimulation, mirroring the kinetics of mRNA accumulation (Figure 7B-F; see Figure 6A, B, D, E). In the presence of F14, the inducible recruitment of BRD4 to the *NFKBIA* and *CXCL8* promoters remained unaffected, whilst its recruitment to *CCL2* and *CXCL10* was blocked (Figure 7C). This strongly suggests that inhibition of acetylation of p65 at K310 by F14 is relayed downstream to the recruitment of BRD4 to the “F14-sensitive” promoters, but not to the “F14-resistant” promoters.

The BRD4 recruitment to *NFKBIA* and *CXCL8* promoters despite inhibition of p65-K310 acetylation prompted investigation of whether other acetyl-lysine residues are recognised. The bromodomain-mediated docking onto acetylated lysine residues is generally accepted as responsible for the recruitment of BRD4 to the chromatin [reviewed by [83, 84]]. For instance, histone 4 acetylated on K5, K8 and K12 (H4K5/K8/K12ac) is responsible for BRD4 recruitment to NF-κB-responsive genes upon lipopolysaccharide stimulation [86]. The recruitment of BRD4 to the *NFKBIA*, *CXCL8*, *CCL2*, and *CXCL10* promoters was tested in the presence of the bromodomain inhibitor JQ1, by chromatin immunoprecipitation and quantitative PCR. BRD4 was still recruited to *NFKBIA* and *CXCL8* promoters after TNF-α stimulation in the presence of JQ1, whilst inducible recruitment to *CCL2* and *CXCL10* promoters was abolished by JQ1 (Figure 7H-K). As a control for JQ1 pharmacological activity, BRD4 recruitment to the *CCND1* gene promoter was diminished by this small-molecule inhibitor (Figure S7). *CCND1* is a BRD4 target gene that encodes the cell cycle regulator cyclin D1 and was used as a positive control [87]. Altogether, these results suggest that the inducible recruitment of BRD4 to some promoters is independent of the bromodomains.

### F14 is unique among known viral antagonists of NF-κB

The TAD domain of p65 belongs to the class of acidic activation domains, characterised by a preponderance of aspartic acid or glutamic acid residues surrounding hydrophobic motifs [22]. VP16 is a transcriptional activator from HSV-1 that bears a prototypical acidic TAD (Figure 8A) and inhibits the expression of virus-induced IFN-β by association with p65 and IRF3 [44]. Although the VP16-mediated inhibition of the IFN-β promoter was independent of its TAD, we revisited this observation to investigate the effect of VP16 more specifically on NF-κB-dependent gene activation. VP16 inhibited NF-κB reporter gene expression in a dose-dependent manner and deletion of the TAD reduced NF-κB inhibitory activity of VP16 about 2-fold, but some activity remained (Figure 8A).

**Figure 8:**
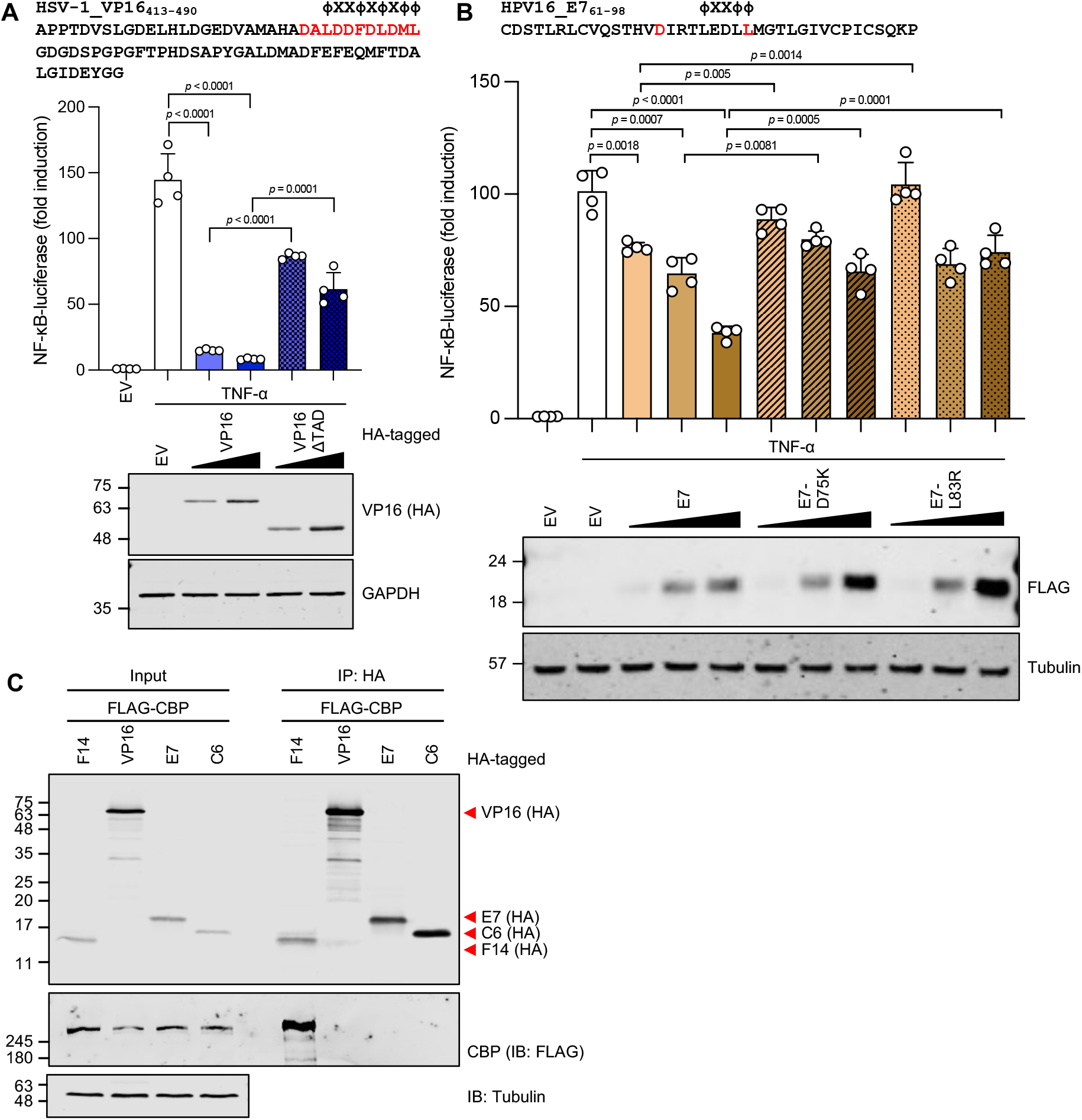
F14 is unique among known viral inhibitors of NF-κB. (**A**) Top: Amino acid sequence of the TAD of HSV-1 VP16 with the acidic activation domain similar to p65 highlighted in red, and hydrophobic residues (Φ) are indicated. Middle: NF-κB-dependent luciferase activity in HEK 293T cells transfected with vectors expressing VP16, VP16 mutant, or empty vector (EV), and stimulated with TNF-α. Bottom: Immunoblotting. (**B**) Top: Amino acid residues 61-98 from HPV16 protein E7 encompassing a ΦXXΦΦ motif containing and preceded by negatively charged residues. Highlighted are two residues mutated to disrupt this motif. Middle: NF-κB-dependent luciferase activity in HEK 293T cells expressing E7 and two mutants as described in (**A**). Bottom: Immunoblotting. Means + s.d. (*n* = 4 per condition) are shown. (**C**) Lysates from transfected HEK 293T cells were immunoprecipitated with anti-HA. Immunoblots are representative of two independent experiments. Protein molecular masses in kDa are shown on the left of the blots. Immunoblots of tagged proteins are labelled with the protein name followed the epitope tag antiboby in parentheses. When multiple tagged proteins are shown in the same immunoblot, each protein is indicated a red arrowhead. Statistical significance was determined by the Student’s *t*-test.

A search for other viral proteins that contain motifs resembling the ΦXXΦΦ motif present in acidic transactivation activation domains detected a divergent ΦXXΦΦ motif in protein E7 (aa 79-83) from HPV16, with acidic residues upstream (D75) or within (E80 and D81) the motif (Figure 8B). E7 has been reported to inhibit NF-κB activation, in addition to its role in promoting cell cycle progression [43, 88–90]. We confirmed that HPV16 protein E7 inhibits NF-κB- dependent gene expression (Figure 8B). Furthermore, E7 mutants harbouring aa substitutions that inverted the charge of D75 (D75K) or added a positive charge to the otherwise hydrophobic L83 (L83R) were impaired in their capacity to inhibit NF-κB (Figure 8B).

Lastly, the ability of VP16 and E7 to associate with CBP was assessed after ectopic expression in HEK 293T cells. Neither VP16 nor E7, like VACV protein C6 used as negative control, co-precipitated CBP under conditions in which F14 did (Figure 8C). These findings indicate that the mimicry of p65 TAD by F14 is a strategy unique among human pathogenic viruses to suppress the activation of NF-κB.

## DISCUSSION

The inducible transcription of NF-κB-dependent genes is a critical response to virus infection. After binding to κB sites in the genome, NF-κB promotes the recruitment of chromatin remodelling factors, histone-modifying enzymes, and components of the transcription machinery. Thereby, NF-κB couples the sensing of viral and inflammatory signals to the selective activation of the target genes. In response, viruses have evolved multiple immune evasion strategies, including interference with NF-κB activation. VACV is a paradigm in viral evasion mechanisms, inasmuch as this poxvirus encodes 15 proteins known to intercept NF- κB activation downstream of PRRs and cytokine receptors [reviewed by [55, 56], [91, 92]]. Nonetheless, a VACV strain lacking all these inhibitors still prevented NF-κB activation after p65 translocation into the nucleus [57], indicating the existence of other inhibitor(s).

Here, VACV protein F14, which is conserved in all orthopoxviruses, including ancient variola viruses, is shown to inhibit NF-κB activation within the nucleus and its mechanism of action is elucidated. First, ectopic expression of F14 reduces NF-κB-dependent gene expression stimulated by TNF-α or IL-1β (Figure 1A, B). Second, F14 is expressed early during VACV infection, and is small enough (8 kDa) to diffuse passively into the nucleus (Figure 1E, H). Third, a VACV strain lacking both A49 and F14 (vv811ΔA49ΔF14) is less able to suppress cytokine-stimulated NF-κB-dependent gene expression than vv811ΔA49 (Figure 1I). Fourth, following TNF-α stimulation, IκBα degradation, IKK-mediated phosphorylation of p65 at S536 and p65 accumulation in the nucleus remained unaffected in the presence of F14 (Figure 2A, B). Lastly, F14 blocked NF-κB-dependent gene expression stimulated by p65 overexpression, indicating that it acts at or downstream of p65 (Figure 2C). Mechanistically, F14 inhibits NF- κB via a C-terminal 23 aa motif that resembles the acidic activation domain of p65. F14 disrupts the binding of p65 to its coactivator CBP (Figures 4 and 5) and reduces acetylation of p65 K310. Subsequently, F14 inhibits the inducible recruitment of BRD4 to *CCL2* and *CXCL10* promoters, but not to *NFKBIA* and *CXCL8* promoters (Figure 7). These findings correlated with F14 suppressing *CCL2* and *CXCL10*, but not *NFKBIA* and *CXCL8*, mRNA expression (Figure 6A, B, D, E). The selective inhibition of a subset of NF-κB-dependent genes by F14, despite the interference with molecular events deemed important for p65-mediated transactivation, underscores the complexity of the nuclear actions of NF-κB. Initial understanding of NF-κB-mediated gene activation was derived mostly using artificial reporter plasmids, but subsequent genome-wide, high-throughput studies uncovered diverse mechanisms of gene activation [14, 16, 24, 28, 29, 86, 93]. Because multiple promoters containing κB sites are preloaded with CBP/p300, RNAP II and general transcription factors, the activation of transcription by NF-κB relies on the recruitment of BRD4 [86, 93].

Recruitment of BRD4 to promoters and enhancers occurs via bromodomain-mediated docking onto acetyl-lysine residues on either histones or non-histone proteins and promotes chromatin remodelling and transcription [reviewed by [83, 84]]. For NF-κB-bound promoters, BRD4 recognises p65 acetylated at K310 [39]. This explains how F14 reduced inducible enrichment of BRD4 on the *CCL2* and *CXCL10* promoters following TNF-α stimulation: namely, reduced acetylation of p65 by CBP (Figure 7A, B, E, F). Nonetheless, BRD4 enrichment on the *NFKBIA* and *CXCL8* promoters remained unaffected in the presence of F14 (Figure 7C, D). Genes whose expression is resistant to F14 inhibition might be activated independently of the p65 TA1 domain, as is the case for some NF-κB-responsive genes in mouse fibroblasts stimulated with TNF-α, including *Nfkbia*. For those genes, p65 occupancy on the promoter elements suffices for gene activation, via recruitment of secondary transcription factors [24]. However, BRD4 enrichment on *NFKBIA* and *CXCL8* promoters also remained unaffected in the presence of the bromodomain inhibitor JQ1 (Figure 7H, I), indicating alternative mechanism(s) of BRD4 recruitment to some promoters. Downstream of p65, alternative recruitment via protein-protein interactions through the C-terminal domains of BRD4 might mediate BRD4 recruitment to some NF-κB-bound promoters independently of the recognition of acetyl-lysine residues by the N-terminal bromodomains, which is recognised as the main mechanism of BRD4 recruitment to chromatin [reviewed by [83, 84]]. For instance, BRD4 interacts directly with multiple transcription factors and chromatin remodellers independently of acetylation [94]. Further investigation of acetylation-independent recruitment of BRD4 to inducible promoters observed here and elsewhere [95] is warranted.

In the nucleus, p65 engages with multiple binding partners via its transactivation domains, including the direct interactions between TA1 and TA2 and the KIX and transcriptional adaptor zinc finger (TAZ) 1 domains of CBP, respectively. These interactions are mediated by hydrophobic contacts of the ΦXXΦΦ motifs and complemented by electrostatic contacts by the acidic residues in the vicinity of the hydrophobic motifs [79, 93]. Sequence analysis suggested that F14 mimics the p65 TA1 domain (Figure 3A). Indeed, fusion of the TAD-like domain of F14 to a p65 mutant lacking the TA1 domain restored its transactivation activity to wildtype levels (Figure 3E). This explains the observation from a yeast two-hybrid screen of VACV protein-protein interactions, in which F14 could not be tested because it was found to be a strong activator when fused to the GAL4 DNA-binding domain [96]. Site-directed mutagenesis of F14 revealed that the dipeptide D62/63, but not L65 or L68 of the ΦXXΦΦ motif, is required for inhibition of NF-κB (Figure 4A), for interaction with CBP (Figure 4B) and for the efficient disruption of p65 binding to CBP (Figure 5D). This contrasts with the molecular determinants of p65 TA1 function, i.e., both hydrophobic (F542) and acidic (including D539 and D541) residues contribute to p65 TA1 transactivation activity [80, 81]. Although the p65 TA1 binding to the KIX domain of CBP was shown to depend on F542, the importance of the electrostatic interactions by D539 and D541 is yet to be tested [79]. Of note, a recent high-throughput mutagenesis analysis of a model acidic activation domain provided useful insight into the relative contributions of hydrophobic and acidic residues for transcriptional activity. This analysis supports a model in which key hydrophobic residues require the acidic residues to keep them exposed to solvent where they can interact with coactivators [97]. We cannot rule out that F14 function depends on other C-terminal hydrophobic residues, but our observation that F14-D62/63K (F14-D62 aligns with p65-D539) mutant is impaired in disrupting p65-KIX interaction in cells is in line with the hypothesis of how acidic activation domains work. Future elucidation of the structure of F14-KIX complex and its comparison with p65 TA1-KIX co-structure will be necessary to address this apparent discrepancy. The “imperfect” nature of F14 mimicry is not without precedent in poxviruses. VACV protein A49, the mimic of IκBα phosphodegron, contains an extra aa residue between the two phosphorylatable serine residues of the degron and requires the phosphorylation of just one of the two serines to interact with the E3 ligase β-TrCP and thus to prevent IκBα degradation [54].

The diminished acetylation of p65 K310 is a direct consequence of the disruption of CBP and p65 interaction by F14. Other poxvirus proteins are reported to inhibit p65 acetylation. For instance, ectopic expression of VACV protein K1 inhibited CBP-dependent p65 acetylation and NF-κB-dependent gene expression [98], whilst during infection, K1 inhibited NF-κB activation upstream of IκBα degradation [99]. Regardless of whether K1 inhibits NF-κB upstream or downstream of p65, the vv811ΔA49 strain used to predict the existence of additional VACV inhibitors of NF-κB lacks K1 [57]. The other poxviral protein that inhibits CBP-mediated acetylation of p65, and thereby NF-κB activation, is encoded by gene *002* of orf virus, a parapoxvirus that causes mucocutaneous infections in goats and sheep [100]. However, protein 002 differs from F14 in that it interacts with p65 to prevent phosphorylation at p65-S276 and the subsequent acetylation at K310 by p300 [100, 101].

This study adds VACV protein F14 to the list of viral binding partners of CBP and its paralogue p300, which includes adenovirus E1A protein [102], human immunodeficiency virus (HIV) 1 Tat protein [103], human T-cell lymphotropic virus (HTLV) 1 Tax protein [104], high-risk HPV16 E6 protein [105], and polyomavirus T antigen [106]. Despite the fact that some of these proteins also inhibit NF-κB activation [43, 105, 107], F14 is unique among them in mimicking p65 TA1 to bind to CBP and prevent its interaction with p65. HPV16 E6 also disrupts the interaction of CBP with p65 but, unlike F14, E6 lacks a ΦXXΦΦ motif surrounded by acidic residues and inhibits the expression of CXCL8 and therefore is mechanistically distinct [43]. After searching for additional viral proteins that might mimic p65 TAD, we focused on HPV16 E7 and HSV-1 VP16. The latter protein has a prototypical acidic TAD (Figure 8A), the former bears a motif resembling the ΦXXΦΦ motif (Figure 8B), and both proteins inhibit NF-κB activation [43, 44, 88–90]. Data presented here confirm that VP16 and E7 each inhibit NF-κB- dependent gene expression (Figure 8A, B). However, neither co-precipitated CBP under conditions in which F14 did (Figure 8C), suggesting VP16 and E7 inhibit NF-κB activation by a mechanism distinct from F14. The interaction between VP16 and CBP is contentious [21, 44] and data presented here suggest that these two proteins do not associate with each other under the conditions tested. Therefore, the molecular mimicry of F14 might be only rivalled by that of the avian reticuloendotheliosis virus, a retrovirus whose *v-Rel* gene was acquired from an avian host. A viral orthologue of c-Rel with weak transcriptional activity, v-Rel acts as dominant-negative protein to repress NF-κB-dependent gene activation in avian cells [108].

Overall, our search for additional inhibitors of NF-κB activation encoded by VACV unveiled a viral strategy to inhibit this transcription factor that is unique among known viral antagonists of NF-κB. By mimicking the TA1 domain of p65, F14 disrupts the interaction between p65 and its coactivator CBP, thus inhibiting the downstream molecular events that trigger the activation of a subset of inflammatory genes in response to cytokine stimulation. Among these events, the recruitment of RNAP II processivity factor BRD4 is important for induction of the inflammatory response. This study also showed BRD4 is recruited to some inducible NF-κB-dependent promoters independently of the recognition of acetylated chromatin (i.e., acetyl-lysine residues), via an unknown mechanism that warrants further investigation. Two lines of evidence illustrated the biological importance of F14. First, F14-D62/63 site is conserved in F14 orthologues from different orthopoxviruses, including human pathogens cowpox and monkeypox viruses, and ancient (10^th^ century CE) and modern variola virus strains (Figure 3A). Second, a VACV strain lacking F14 is attenuated in an intradermal model of infection (Figure 1F), despite the presence of several other VACV-encoded NF-κB inhibitors [reviewed by [55, 56]]. The attenuation of vΔF14 also shows the function of F14 is not redundant with these other inhibitors of NF-κB, despite the selective inhibition imparted by F14 (Figure 6A-F). From the viral perspective, the selective inhibition of only a subset of NF-κB-responsive genes by F14 might represent an adaptation to counteract the host immune response more efficiently. If an NF-κB-activating signal reached the nucleus of an infected cell, maintaining expression of some NF-κB-dependent genes, particularly *NFKBIA*, might promote the signal termination by IκBα. Newly synthesised IκBα not only tethers cytoplasmic NF-κB, but can also remove NF-κB from the DNA and cause its export from the nucleus [17, 18, 109]. We anticipate that other viruses might also use the selective inhibition of NF-κB to exploit the pro-viral functions of active NF-κB whilst dampening its pro-inflammatory and antiviral activities.

## ACKNOWLEDGEMENTS

The authors thank Rachel Seear, Stephanie Macilwee, and Jemma Milburn for technical support, and Florian Pfaff and Martin Beer (Friedrich-Loeffler-Institut, Germany) for help with access to cowpox RNA sequencing dataset. We also thank John Doorbar (Dept. Pathology, University of Cambridge, UK), Colin Crump (Dept. Pathology, University of Cambridge, UK), Tony Kouzarides (Dept. Pathology and The Gurdon Institute, University of Cambridge, UK), and Gerd Blobel (University of Pennsylvania, Philadelphia, USA) for providing us with reagents. We are also grateful to Tony Kouzarides for helpful advice and to Callum Talbot-Cooper for critical reading of the manuscript.

## AUTHOR CONTRIBUTION

Conceptualisation: JDA, AAT, GLS

Methodology: JDA, HR, AAT, EVS, CAM, AJB, MPB, BYWC

Software: N/A

Validation: JDA, HR, AAT, EVS

Formal Analysis: JDA, HR

Investigation: JDA, HR, AAT, EVS

Resources: AAT, CAM, AJB, BYWC, GLS

Data Curation: JDA

Writing – Original Draft Preparation: JDA

Writing – Review and Editing: JDA, HR, AAT, CAM, AJB, BYWC, GLS

Visualisation: JDA

Supervision: JDA, GLS

Project Administration: JDA, GLS

Funding: JDA, GLS

## MATERIAL AND METHODS

### Sequence analysis

Candidate open reading frames (ORFs) encoding the unknown VACV inhibitor of NF-κB were first selected based on VACV genomes available on the NCBI database (accession numbers: NC_006998.1 for the Western Reserve strain, and M35027.1 for the Copenhagen strain). The prediction of molecular mass and isoelectric point (pI), and of nuclear localisation signal (NLS) sequences, of the candidate VACV gene products was done with ExPASy Compute pI/MW tool (https://web.expasy.org/compute_pi/) and SeqNLS (http://mleg.cse.sc.edu/seqNLS/), [110], respectively. Domain searches were performed using InterPro (http://www.ebi.ac.uk/interpro/search/sequence/), UniProt (https://www.uniprot.org/uniprot/), HHpred (https://toolkit.tuebingen.mpg.de/tools/hhpred), PCOILS (https://toolkit.tuebingen.mpg.de/tools/pcoils), and Phobius (https://www.ebi.ac.uk/Tools/pfa/phobius/) [111]. Gene family searches were done within the Pfam database (https://pfam.xfam.org/) and conservation within the poxvirus family, with Viral Orthologous Clusters (https://4virology.net/virology-ca-tools/vocs/) [74] and protein BLAST (https://blast.ncbi.nlm.nih.gov/Blast.cgi) searches. Phyre2 (http://www.sbg.bio.ic.ac.uk/phyre2/html/page.cgi?id=index) [78] was used for the prediction of F14 protein structure. Multiple sequence alignments were performed using Clustal Omega (https://www.ebi.ac.uk/Tools/msa/clustalo/) and ESPript 3.0 (http://espript.ibcp.fr/ESPript/ESPript/) [112] was used for the visualisation of protein sequence alignments. All poxvirus sequences referred to in this study are listed in Table S1.

### Expression vectors

The VACV *F6L*, *F7L*, *F14L*, *A47L*, *B6R*, *B11R*, and *B12R* ORFs (strains Western Reserve and Copenhagen, if sequences diverged between strains) were codon-optimised for expression in human cells and synthesised by GeneArt (Thermo Fisher Scientific), with an optimal 5’ Kozak sequence and fused to an N-terminal FLAG epitope. For ease of subsequent subcloning, 5’ *Bam*HI and 3’ *Xba*I restriction sites were included as well as a *Not*I site + 1G between the epitope tag and the ORF. The *Not*I site + 1G generates an (Ala)3 linker between the epitope tag and the protein of interest. For mammalian expression, nucleotide sequences encoding N-terminal FLAG-tagged VACV proteins were subcloned between the *Bam*HI and *Xba*I restriction sites of a pcDNA4/TO vector (Invitrogen). Alternatively, codon-optimised F14 was PCR-amplified to include a 3’ HA tag or a 3’ FLAG or no epitope tag, and 5’ *Bam*HI and 3’ *Xba*I sites to clone into pcDNA4/TO plasmid. In addition, codon-optimised F14 sequence was PCR-amplified to include 5’ *Bam*HI and 3’ *Not*I sites to facilitate cloning into a pcDNA4/TO-based vector containing a TAP tag sequence after the *Not*I site; the TAP tag consisted of two copies of the Strep-tag II epitope and one copy of the FLAG epitope [113]. Mutant F14 expression vectors were constructed with QuikChange II XL Site-Directed Mutagenesis kit (Agilent), using primers containing the desired mutations and C-terminal TAP-tagged codon-optimised F14 cloned into pcDNA4/TO as template. The pcDNA4/TO-based expression vectors for VACV proteins C6 and B14 have been described [66, 114].

The ORF encoding HPV16 E7 protein was amplified from a template kindly provided by Dr. Christian Kranjec and Prof. John Doorbar (Dept. Pathology, Cambridge, UK) and cloned into 5’ *Bam*HI and 3’ *Not*I sites of a pcDNA4/TO-based vectors fused to a C-terminal TAP tag or HA epitope. Vectors expressing mutant E7 proteins were generated by site-directed mutagenesis as described above. The ORF encoding HSV-1 VP16 and ΔTAD mutant (lacking aa 413-490) were amplified from a pEGFP-C2-based VP16 expression plasmid kindly provided by Dr. Colin Crump (Dept. Pathology, Cambridge, UK) and cloned into 5’ *Bam*HI and 3’ *Not*I sites of a pcDNA4/TO-based vector fused to a C-terminal HA epitope.

The pcDNA4/TO plasmids encoding TAP- and HA-tagged p65 were described elsewhere [91], and plasmids expressing FLAG-tagged mouse CBP (pCMV5-CBP-FLAG), and HA-tagged mouse CBP (pRcRSV-CBP-HA) were kind gifts from Prof. Gerd A. Blobel (University of Pennsylvania, Philadelphia, USA) and Prof. Tony Kouzarides (Dept. Pathology and The Gurdon Institute, Cambridge, UK), respectively. Firefly luciferase reporter plasmids for NF-κB, STAT and AP-1, as well as the constitutively active TK-*Renilla* luciferase reporter plasmid were kind gifts from Prof. Andrew Bowie (Trinity College, Dublin, Republic of Ireland). The NF- κB, AP-1 and STAT reporter plasmids encode firefly luciferase under the control of consensus NF-κB response element repeats [(GGGAATTTCC)5], AP-1 response element repeats [(TGACTAA)7] and IFN-stimulated response element (ISRE) [(TAGTTTCACTTTCCC)5], respectively.

The oligonucleotide primers used for cloning and site-directed mutagenesis are listed in Table S2. Nucleotide sequences of the inserts in all the plasmids were verified by Sanger DNA sequencing.

### Cell lines

All cell lines were grown in medium supplemented with 10% foetal bovine serum (FBS, Pan Biotech), 100 units/mL of penicillin and 100 µg/mL of streptomycin (Gibco), at 37⁰C in a humid atmosphere containing 5% CO2. Human embryo kidney (HEK) 293T epithelial cells (ATCC, CRL-11268), and monkey kidney BS-C-1 (ATCC, CCL-26) and CV-1 (ATCC, CCL-70) epithelial cells were grown in Dulbecco’s modified Eagle’s medium (DMEM, Gibco). Rabbit kidney RK13 epithelial cells (ATCC, CCL-37) were grown in minimum essential medium (MEM, Gibco) and human cervix HeLa epithelial cells (ATCC, CCL-2), in MEM supplemented with non-essential amino acids (Gibco). T-REx-293 cells (Invitrogen) were grown in DMEM supplemented with blasticidin (10 µg/mL, InvivoGen), whilst the growth medium of T-REx-293- derived cell lines stably transfected with pcDNA4/TO-based plasmids was further supplemented with zeocin (100 µg/mL, Gibco).

The absence of mycoplasma contamination in the cell cultures was tested routinely with MycoAlert detection kit (Lonza), following the manufacturer’s recommendations.

### Construction of recombinant viruses

A VACV Western Reserve (WR) strain lacking F14 (vΔF14) was constructed by introduction of a 137-bp internal deletion in the *F14L* ORF by transient dominant selection [115]. A DNA fragment including the first 3 bp of *F14L* ORF and 297 bp upstream, intervening *NotI* and *Hind*III sites, and the last 82 bp of the ORF and 218 bp downstream were generated by overlapping PCR and inserted into the *Pst*I and *Bam*HI sites of pUC13-Ecogpt-EGFP plasmid, containing the *Escherichia coli* guanylphosphoribosyl transferase (Ecogpt) gene fused in-frame with the enhanced green fluorescent protein (EGFP) gene under the control of the VACV 7.5K promoter [70]. The resulting plasmid contained an internal deletion of the *F14L* ORF (nucleotide positions 42049-42185 from VACV WR reference genome, accession number NC_006998.1). The remaining sequence of *F14L* was out-of-frame and contained multiple stop codons, precluding the expression of a truncated version of F14. The derived plasmid was transfected into CV-1 cells that had been infected with VACV-WR at 0.1 p.f.u./cell for 1 h. After 48 h, progeny viruses that incorporated the plasmid by recombination and expressed the Ecogpt-EGFP were selected and plaque-purified three times on monolayers of BS-C-1 cells in the presence of mycophenolic acid (25 μg/mL), supplemented with hypoxanthine (15 μg/mL) and xanthine (250 μg/mL). The intermediate recombinant virus was submitted to three additional rounds of plaque purification in the absence of the selecting drugs and GFP-negative plaques were selected. Under these conditions, progeny viruses can undergo a second recombination that result in loss of the Ecogpt-EGFP cassette concomitantly with either incorporation of the desired mutation (vΔF14) or reversal to wildtype genotype (vF14). Because vΔF14 and vF14 are sibling strains derived from the same intermediate virus, they are genetically identical except for the 137-bp deletion in the *F14L* locus. Viruses were analysed by PCR to identify recombinants by distinguishing wildtype and ΔF14 alleles, and the presence or absence of the Ecogpt-EGFP cassette.

To restore F14 expression in vΔF14, the *F14L* locus was amplified by PCR, including about 250 bp upstream and downstream of the ORF, and inserted into the *Pst*I and *Bam*HI sites of pUC13-Ecogpt-EGFP plasmid. Additionally, *F14L* ORF fused to the sequence coding a C-terminal TAP tag was also amplified by overlapping PCR, including the same flanking sequences described above, and inserted into the *Pst*I and *Bam*HI sites of pUC13-Ecogpt- EGFP plasmid. By using the same transient dominant selection method, these plasmids were used to generate two revertant strains derived from vΔF14: (i) vF14-Rev, in which F14 expression from its natural locus was restored, and (ii) vF14-TAP, expressing F14 fused to a C-terminal TAP tag under the control of its natural promoter. The vC6-TAP virus was described elsewhere [114].

A VACV vv811 strain lacking both A49 and F14 (vv811ΔA49ΔF14) was also constructed by transient dominant selection. The resultant virus contained the same 137-bp internal deletion in the *F14L* ORF within the vv811ΔA49 strain generated previously [57]. The distinction between wildtype and ΔF14 alleles in the obtained viruses, and the presence or absence of the Ecogpt-EGFP cassette, was determined by PCR analysis.

The oligonucleotide primers used to generate the recombinant VACV strains are listed in Table S2. To verify that all the final recombinant viruses harboured the correct sequences, PCR fragments spanning the *F14L* locus were sequenced.

### Preparation of virus stocks

The stocks of virus strains derived from VACV WR were prepared in RK13 cells. Cells grown to confluence in T-175 flasks were infected at 0.01 p.f.u./cell until complete cytopathic effect was visible. The cells were harvested by centrifugation, suspended in a small volume of DMEM supplemented with 2% FBS, and submitted to multiple cycles of freezing/thawing and sonication to lyse the cells and disrupt aggregates of virus particles and cellular debris. These crude virus stocks were used for experiments in cultured cells. Crude stocks of vv811 and derived strains were prepared in the same way, except for the BS-C-1 cells used for infection. For the *in vivo* work, virus stocks were prepared by ultracentrifugation of the cytoplasmic fraction of infected cell lysates through sucrose cushion and suspension of the virus samples in 10 mM Tris-HCl pH 9.0 [116]. The viral titres in the stocks were determined by plaque assay on BS-C-1 cells.

### Virus growth and spread assays

To analyse virus growth properties in cell culture, single-step growth curve experiments were performed in HeLa cells. Cells were grown to about 90% confluence in T-25 flasks and then infected at 5 p.f.u./cell in growth medium supplemented with 2% FBS. Virus adsorption was at 37⁰C for 1 h. Then the inoculum was removed, and the cells were replenished with growth medium supplemented with 2% FBS. At 1, 8, and 24 h p.i., infected-cell supernatants and monolayers were collected for determination of extracellular and cell-associated infectious virus titres, respectively, by plaque assay on BS-C-1 cells. Supernatants were clarified by centrifugation to remove cellular debris and detached cells, whereas cell monolayers were scraped and disrupted by three cycles of freezing/thawing followed by sonication, to release intracellular virus particles.

The virus spread in cell culture was assessed by plaque formation. Confluent monolayers of BS-C-1 cells in 6-well plates were infected with 50 p.f.u./well and overlaid with MEM supplemented with 2% FBS and 1.5% carboxymethylcellulose. After 48 h, infected-cell monolayers were stained with 0.5% crystal violet solution in 20% methanol and imaged.

### Construction of inducible F14-expressing T-REx-293 cell line

T-REx-293 cells (Invitrogen), which constitutively expresses the Tet repressor (TetR) under the control of the human cytomegalovirus (HCMV) immediate early promoter, were transfected with pcDNA4/TO-coF14-TAP plasmid, which encodes human codon-optimised F14 fused to a C-terminal TAP tag under the control of the HCMV immediate early promoter and two tetracycline operator 2 (TetO2) sites. Transfected cells were selected in the presence of blasticidin (10 µg/mL) and zeocin (100 µg/mL) and clonal cell lines were obtained by limiting dilution. Expression of protein F14 within these clones was analysed by immunoblotting and flow cytometry with anti-FLAG antibodies. T-REx-293-EV, T-REx-293-B14, and T-REx-293- C6 cell lines were described elsewhere [63].

### Reporter gene assays

HEK 293T cells in 96-well plates were transfected in quadruplicate with firefly luciferase reporter plasmid (NF-κB, ISRE, or AP-1), TK-*Renilla* luciferase reporter plasmid (as an internal control) and the desired expression vectors or empty vector (EV) using *Trans*IT-LT1 transfection reagent (Mirus Bio), according to the manufacturer’s instruction. On the following day, cells were stimulated with TNF-α (10 ng/ml, PeproTech) or IL-1β (20 ng/ml, PeproTech) for 8 h (for NF-κB activation), IFN-α2 (1000 U/ml, PeproTech) for 8 h (for IFNAR1/STAT activation), or phorbol 12-myristate 13-acetate (10 ng/ml) for 24 h (for MAPK/AP-1 activation). Alternatively, NF-κB was activated by co-transfection of p65-overexpressing plasmid and cells were harvested 24 h after transfection. To test the effect of increasing amounts of viral proteins, five-fold dilutions of the desired expression vectors were used (5 and 25 ng, or 5, 25, and 125 ng, depending on the experiment). The total amount of transfected DNA was made equivalent by addition of empty vector.

To measure NF-κB-luciferase activation during infection, A549 cells transduced with a lentiviral vector expressing the firefly luciferase under the control of an NF-κB promoter (A549- NF-κB-Luc) [57] were grown in 96-well plates and infected with VACV vv811 and derived strains at 5 p.f.u./cell. After 6 h, cells were stimulated with TNF-α (10 ng/ml, PeproTech) or IL- 1β (20 ng/ml, PeproTech) for an additional 6 h. In parallel, A549-NF-κB-Luc cells grown in 6- well plates were infected with the equivalent amount of virus for 12 h and cell lysates were analysed by immunoblotting.

Cells were lysed using passive lysis buffer (Promega) and firefly and *Renilla* luciferase luminescence was measured using a FLUOstar luminometer (BMG). During the measurement, the gain was adjusted to keep the reads within the dynamic range of the luminometer. Home-made substrates for firefly luciferase (20 mM tricine, 2.67 mM MgSO4.7H2O, 0.1 mM EDTA, 33.3 mM DTT, 0.53 mM ATP, 0.27 mM acetyl coenzyme A, 0.5 mM D-luciferin (Nanolight Technology), 0.3 mM magnesium carbonate hydroxide, pH 7.8] and *Renilla* luciferase [2 μg/ml native coelenterazine (Nanolight Technology) in PBS] were used. Promoter activity was obtained by calculation of firefly/*Renilla* luciferase ratios and the promoter activity under pathway stimulation was normalised to the activity of the respective non-stimulated control of each protein under test. In parallel, aliquots of the replicas of each condition tested were combined, mixed with 5 × SDS-gel loading buffer, and immunoblotted to confirm the expression of the proteins tested.

### Virus infection in cell culture

HeLa or HEK 293T cells in 6-well plates (for protein expression analyses) or 10-cm dishes (for immunoprecipitation experiments) were infected at 5 p.f.u./cell. Viral inocula were prepared in growth medium supplemented with 2% FBS. Viral adsorption was done at 37⁰C for 1 h, after which the medium supplemented with 2% FBS was topped up to the appropriate vessel volume and cells were incubated at 37⁰C.

### In vivo experiments

All animal experiments were conducted according to the Animals (Scientific Procedures) Act 1986 under the license PPL 70/8524. Mice were purchased from Envigo and housed in specific pathogen-free conditions in the Cambridge University Biomedical Services facility.

For the intradermal (i.d.) model of infection, female C57BL/6 mice (6-8-week old) were inoculated with 10^4^ p.f.u. in both ear pinnae and the diameter of the lesion was measured daily using a calliper [117]. For the intranasal (i.n.) model, female BALB/c mice (6-8-week old) were inoculated 5 × 10^3^ p.f.u. divided equally into each nostril and were weighed daily [118]. In both cases, viral inocula were prepared in phosphate-buffered saline (PBS) supplemented with 0.01% bovine serum albumin (BSA, Sigma-Aldrich) and the infectious titres in the administered inocula were confirmed by plaque assay.

For quantification of virus replication after the i.d. infection, infected mice were culled 3, 7, and 10 d p.i. and ear tissues were collected, ground in a tissue homogeniser and passed through a 70-μm nylon mesh using DMEM containing 10% FBS. Samples were frozen, thawed and sonicated three times, to liberate cell-associated virus particles, and the infectious titres present were determined by plaque assay on BS-C-1 cells.

### Immunoblotting

For analysis of protein expression, cells were washed with PBS and lysed on ice with cell lysis buffer [50 mM Tris-HCl pH 8.0, 150 mM NaCl, 1 mM EDTA, 10% (v/v) glycerol, 1% (v/v) Triton X-100 and 0.05% (v/v) Nonidet P-40 (NP-40)], supplemented with protease (cOmplete Mini, Roche) and phosphatase (PhosSTOP, Roche) inhibitors, for 20 min. Lysed cells were scraped and lysates were clarified to remove insoluble material by centrifugation at 17,000 × g for 15 min at 4⁰C. Protein concentration in the cell lysate was determined using a bicinchoninic acid protein assay kit (Pierce). After mixing with 5 × SDS-gel loading buffer and boiling at 100⁰C for 5 min, equivalent amounts of protein samples (15-50 µg/well) were loaded onto SDS-polyacrylamide gels or NuPAGE 4 to 12% Bis-Tris precast gels (Invitrogen), separated by electrophoresis and transferred onto nitrocellulose membranes (GE Healthcare). Membranes were blocked at room temperature with either 5% (w/v) non-fat milk or 3% (w/v) BSA (Sigma-Aldrich) in Tris-buffered saline (TBS) containing 0.1% (v/v) Tween-20. To detect the expression of the protein under test, the membranes were incubated with specific primary antibodies diluted in blocking buffer at 4⁰C overnight. After washing with TBS containing 0.1% (v/v) Tween-20, membranes were probed with fluorophore-conjugated secondary antibodies (LI-COR Biosciences) diluted in 5% (w/v) non-fat milk at room temperature for 1 h. After washing, membranes were imaged using the Odyssey CLx imaging system (LI-COR Biosciences), according to the manufacturer’s instructions. For quantitative analysis of protein levels, the band intensities on the immunoblots were quantified using the Image Studio software (LI-COR Biosciences). The antibodies used for immunoblotting are listed in Table S3.

### Co-immunoprecipitation and pulldown assays

HEK 293T or HeLa cells in 10-cm dishes were infected at 5 p.f.u./cell for 8 h or transfected overnight with the specified epitope-tagged plasmids using polyethylenimine (PEI, Polysciences, 2 μl of 1 mg/ml stock per μg of plasmid DNA). For the competition assays, cells were starved of FBS for 3 h and stimulated with TNF-α (40 ng/ml, PeproTech) in FBS-free DMEM for 15 min before harvesting. Cells were washed with ice-cold PSB, scraped in immunoprecipitation (IP) buffer [50 mM Tris-HCl pH 7.4, 150 mM NaCl, 0.5% (v/v) NP-40, 0.1 mM EDTA], supplemented with protease (cOmplete Mini, Roche) and phosphatase (PhosSTOP, Roche) inhibitors, on ice, transferred to 1.5-ml microcentrifuge tubes and rotated for 30 min at 4⁰C. Cell lysates were centrifuged at 17,000 × g for 15 min at 4⁰C and the soluble fractions were incubated with 20 μl of one of the following affinity resins equilibrated in IP buffer: (i) anti-FLAG M2 agarose (Sigma-Aldrich, Cat# A2220) for IP of FLAG- or TAP-tagged proteins; (ii) anti-HA agarose (Sigma-Aldrich, Cat# A2095) for IP of HA-tagged proteins; or (iii) Strep-Tactin Superflow agarose (IBA, Cat# 2-1206-025) for pulldown of TAP-tagged protein via Strep-tag II epitope. After 2 h of rotation at 4⁰C, the protein-bound resins were washed three times with ice-cold IP buffer. The bound proteins were eluted by incubation with 2× SDS-gel loading buffer and boiled at 100⁰C for 5 min before analysis by SDS-polyacrylamide gel electrophoresis and immunoblotting, along with 10% input samples collected after clarification of cell lysates. The antibodies used for immunoprecipitation are listed in Table S3.

### Reverse transcription and quantitative PCR

To analyse mRNA expression of NF-κB-responsive genes, T-REx-293-F14 in 12-well plates were left uninduced or induced overnight with 100 ng/ml doxycycline (Melford, UK) to induce the expression of F14. Alternatively, T-REx-293-F14 and C6 in 12-well plates were induced overnight with 100 ng/ml doxycycline (Melford, UK) to induce the expression of the VACV proteins. The next day, cells were starved for 3 h by removal of serum from the medium and then stimulated in duplicate with TNF-α (40 ng/ml, PeproTech) in FBS-free DMEM for 0, 1 or 6 h. RNA was extracted using RNeasy Mini Kit (Qiagen) and complementary DNA (cDNA) was synthesised using SuperScript III reverse transcriptase (Invitrogen) and oligo-dT primers (Thermo Scientific), according to the instructions of the respective manufacturers. The mRNA levels of *CCL2*, *CXCL8*, *CXCL10*, *GAPDH* and *NFKBIA* were quantified by quantitative PCR using gene-specific primer sets, fast SYBR green master mix (Applied Biosystems) and the ViiA 7 real-time PCR system (Life Technologies). The oligonucleotide primers used for the qPCR analysis of gene expression are listed in Table S2. Fold-induction of the NF-κB- responsive genes was calculated by the 2^-ΔΔCt^ method using non-induced and non-stimulated T-REx-293-F14 cells, or induced and non-stimulated T-REx-293-C6 cells, as the reference sample, and *GAPDH* as the housekeeping control gene.

### Enzyme-linked immunosorbent assay (ELISA)

The secretion of CXCL8 and CXCL10 was measured by ELISA. T-REx-293-EV, T-REx-293- B14, T-REx-293-C6 and T-REx-293-F14 cells in 12-well plates were incubated overnight in the presence or absence of 100 ng/ml doxycycline (Melford, UK) to induce VACV protein expression. The next day, cells were stimulated in triplicate with TNF-α (40 ng/ml, PeproTech) in DMEM supplemented with 2% FBS for 16 h. To test the effect of the pharmacological inhibition of BRD4 bromodomains, cells were treated with (+)-JQ1 (5 µM, Abcam, Cat# ab141498, dissolved in DMSO) or the equivalent amount of DMSO [0.025% (v/v)] 30 min before TNF-α stimulation. The supernatants were assayed for human CXCL8 and CXCL10 using the respective DuoSet ELISA kits (R&D Biosystems), according to the manufacturer’s instructions.

### Immunofluorescence

For immunofluorescence microscopy, T-REx-293-EV, T-REx-293-B14, T-REx-293-C6 and T-REx-293-F14 cells were grown on poly-D-lysine-treated glass coverslips placed inside 6-well plates. Following induction of protein expression with 100 ng/ml doxycycline (Melford, UK) overnight, cells were starved of FBS for 3 h and then stimulated with 40 ng/ml TNF-α (PeproTech) in FBS-free DMEM for 15 min. At the moment of harvesting, the cells were washed twice with ice-cold PBS and fixed in 4% (v/v) paraformaldehyde for 10 min. After quenching of free formaldehyde with 150 mM ammonium chloride for 5 min, the fixed cells were permeabilised with 0.1% (v/v) Triton X-100 in PBS for 5 min and blocked with 10% (v/v) FBS in PBS for 30 min. Staining was carried out with primary antibodies for 1 h, followed by incubation with the appropriate AlexaFluor fluorophore-conjugated secondary antibodies (Invitrogen Molecular Probes) for 30 min and mounting onto glass slides with Mowiol 4-88 (Calbiochem) containing 0.5 μg/ml DAPI (4’,6-diamidino-2-phenylindole, Biotium). Images were acquired on an LSM 700 confocal microscope (Zeiss) using ZEN system software (Zeiss). Quantification of nuclear localisation of p65 was done manually on the ZEN lite software (blue edition, Zeiss). The details about the antibodies used for immunofluorescence are listed in Table S3.

### Flow cytometry

T-REx-293-F14 cells were induced overnight with 100 ng/ml doxycycline (Melford, UK) in the presence or absence of 10 μM MG132 (Abcam). Cells were detached with trypsin-EDTA (Gibco), washed in PBS and fixed with 4% paraformaldehyde in PBS for 10 min at room temperature with intermittent agitation by vortexing. After centrifugation, fixed cells were suspended in PBS containing 0.1% BSA (Sigma-Aldrich). For intracellular staining of F14- TAP, cells were permeabilised with 0.1% saponin (Sigma-Aldrich) in PBS and stained with the mouse monoclonal antibody against the FLAG tag or isotype control, followed by PE goat anti-mouse IgG (Poly4053, BioLegend). Immunostained cells were fixed again with 1% paraformaldehyde in PBS. Data were acquired with a FACScan/Cytek DxP8-upgraded flow cytometry analyser and analysed with FlowJo software.

### Chromatin immunoprecipitation and quantitative PCR (ChIP-qPCR)

T-REx-F14 cells in 15-cm dishes were incubated overnight in the absence or in the presence of 100 ng/ml doxycycline (Melford, UK) to induce F14 expression. The next day, cells were starved of FBS for 3 h and stimulated with TNF-α (40 ng/ml, PeproTech) in FBS-free DMEM for 0, 1 or 5 h. To test the effect of the pharmacological inhibition of BRD4 bromodomains, T-REx-293-EV cells were treated with (+)-JQ1 (5 µM, Abcam, Cat# ab141498, dissolved in DMSO) or the equivalent amount of DMSO [0.025% (v/v)] 30 min before TNF-α stimulation. Cells were crosslinked with 1% (v/v) formaldehyde added directly to the growth medium. After 10 min at room temperature, crosslinking was stopped by the addition of 0.125 M glycine. Cells were then lysed in 0.2% NP-40, 10 mM Tris-HCl pH 8.0, 10 mM NaCl, supplemented with protease (cOmplete Mini, Roche), phosphatase (PhosSTOP, Roche) and histone deacetylase (10 mM sodium butyrate, Sigma-Aldrich) inhibitors, and nuclei were recovered by centrifugation at 600 × g for 5 min at 4⁰C. To prepare the chromatin, nuclei were lysed in 1% (w/v) SDS, 50 mM Tris-HCl pH 8.0, 10 mM EDTA, plus protease/phosphatase/histone deacetylase inhibitors, and lysates were sonicated in a Bioruptor Pico (Diagenode) to achieve DNA fragments of about 500 bp. After sonication, samples were centrifuged at 3,500 × g for 10 min at 4⁰C and supernatants were diluted four-fold in IP dilution buffer [20 mM Tris-HCl pH 8.0, 150 mM NaCl, 2 mM EDTA, 1% (v/v) Triton X-100, 0.01% (w/v) SDS] supplemented with protease/phosphatase/histone deacetylase inhibitors.

Protein G-conjugated agarose beads (GE Healthcare, Cat# 17-0618-02) equilibrated in IP dilution buffer were used to preclear the chromatin for 1 h at 4⁰C with rotation. Before the immunoprecipitation, 20% of the precleared chromatin was kept as input control. Immunoprecipitation was performed with 8 µg of anti-BRD4 antibody (Cell Signalling Technology, #13440) or anti-GFP (Abcam, #ab290), used as negative IgG control, overnight at 4⁰C with rotation. Protein-DNA immunocomplexes were retrieved by incubation with 60 µl of equilibrated protein G-conjugated agarose beads (GE Healthcare), for 2 h at 4⁰C, followed by centrifugation at 5,000 × g for 2 min at 4⁰C. Immunocomplex-bound beads were then washed: (i) twice with IP wash I [20 mM Tris-HCl pH 8.0, 50 mM NaCl, 2 mM EDTA, 1% (v/v) Triton X-100, 0.1% (w/v) SDS]; (ii) once with IP wash buffer II [10 mM Tris-HCl pH 8.0, 250 mM LiCl, 1 mM EDTA, 1% (v/v) NP-40, 1% (w/v) sodium deoxycholate]; and (iii) twice with TE buffer (10 mM Tris-HCl pH 8.0, 1 mM EDTA). Antibody-bound chromatin was eluted with 1% SDS, 100 mM sodium bicarbonate for 15 min at room temperature. Formaldehyde crosslinks were reversed by incubation overnight at 67⁰C in presence of 1 µg of RNase A and 300 mM NaCl, followed by proteinase K digestion for 2 h at 45⁰C. Co-immunoprecipitated DNA fragments were purified using the QIAquick PCR purification kit (Qiagen) and analysed by quantitative PCR targeting the promoter elements of *NFKBIA*, *CXCL8*, *CCL2*, and *CXCL10* genes. The oligonucleotide primers used for the qPCR analysis of ChIP are listed in Table S2. The primers target regions containing consensus κB sites (5’-GGGRNYYYCC-3′, in which R is a purine, Y is a pyrimidine, and N is any nucleotide), prioritising amplicons overlapping areas with histone modification often observed near active regulatory elements (H3K27ac) according to ENCODE Histone Modification database on UCSC Genome Browser (https://genome.ucsc.edu/index.html). Some primers have been described previously [15, 119, 120].

The ChIP-qPCR data were analysed by the fold enrichment method. Briefly, the signals obtained from the ChIP with each antibody were first normalised to the signals obtained from the corresponding input sample (ΔCt = CtIP – CtInput). Next, the input-normalised signals (ΔCt) were normalised to the corresponding 0 time-point control (i.e. ΔΔCt = ΔCt – ΔCt0). The fold enrichment of each time-point was then calculated with the 2^-ΔΔCt^ formula.

### Statistical analysis

Experimental data are presented as means + s.d., or means ± s.e.m. for *in vivo* results, unless otherwise stated in figure legends. Sample size and number of repeats are indicated in the respective sub-legend, when they apply to specific panels, or in the end, when they apply to all above panels in the figure. Statistical significance was calculated by two-tailed unpaired Student’s *t*-test. In the figures, only *p* < 0.05 values are shown above horizontal brackets indicating the samples being compared. GraphPad Prism software (version 8.4.2) was used for statistical analysis.

### Biological materials

All unique materials are readily available from the corresponding authors upon request. The availability of the antibodies recognising VACV antigens and VACV proteins A49, C6 and D8 is limited.

### Data availability

The authors declare that the main data supporting the findings of this study are available within the article and its supplementary material. Extra data are available from the corresponding authors upon request.

## SUPPLEMENTARY MATERIAL

**Figure S1.**
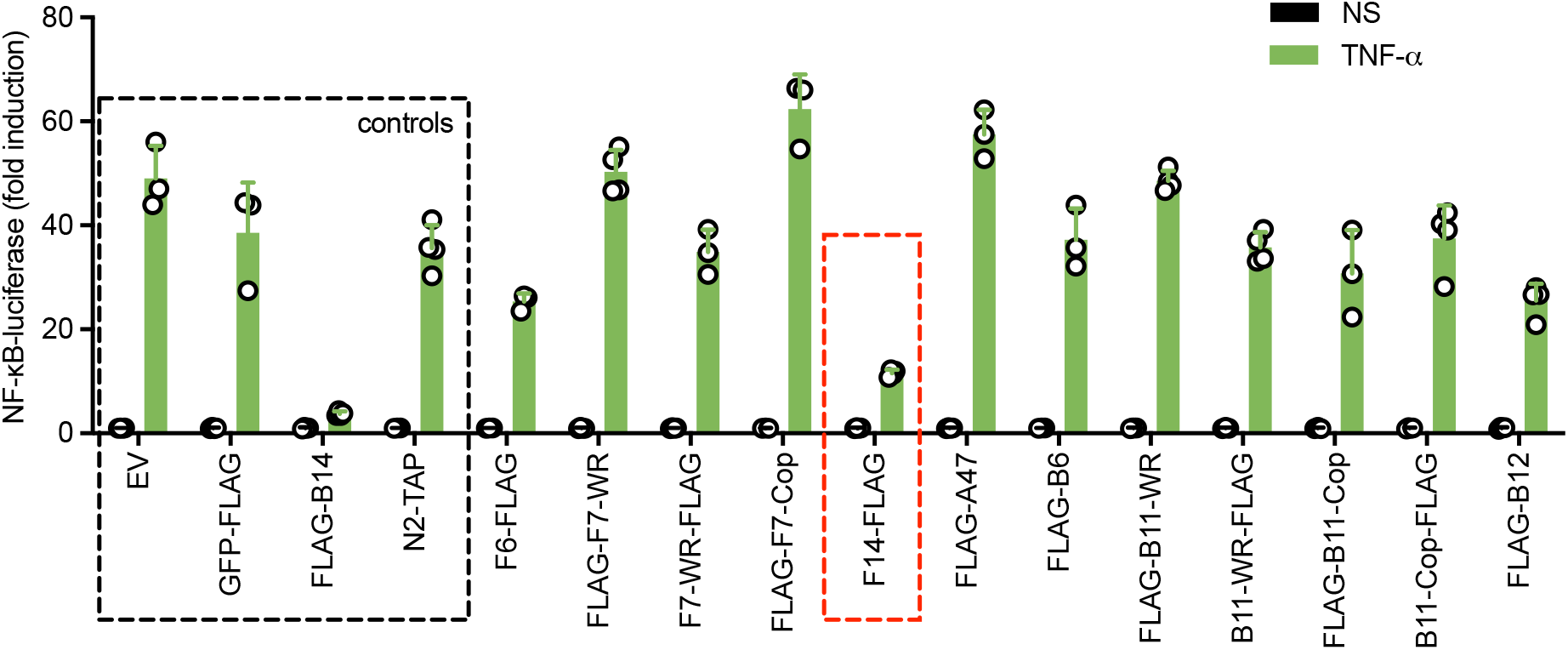
related to Figure 1: Screen of VACV strain WR ORFs for NF-κB inhibitory activity. NF-κB-dependent luciferase activity in HEK 293T cells transfected with vectors expressing the indicated VACV proteins or empty vector (EV), and stimulated with TNF-α. Negative (EV, GFP, and N2) and positive (B14) controls are highlighted in the dashed black square, whilst F14 is highlighted in the dashed red square. Means + s.d. (*n* = 4 per condition) are shown.

**Figure S2.**
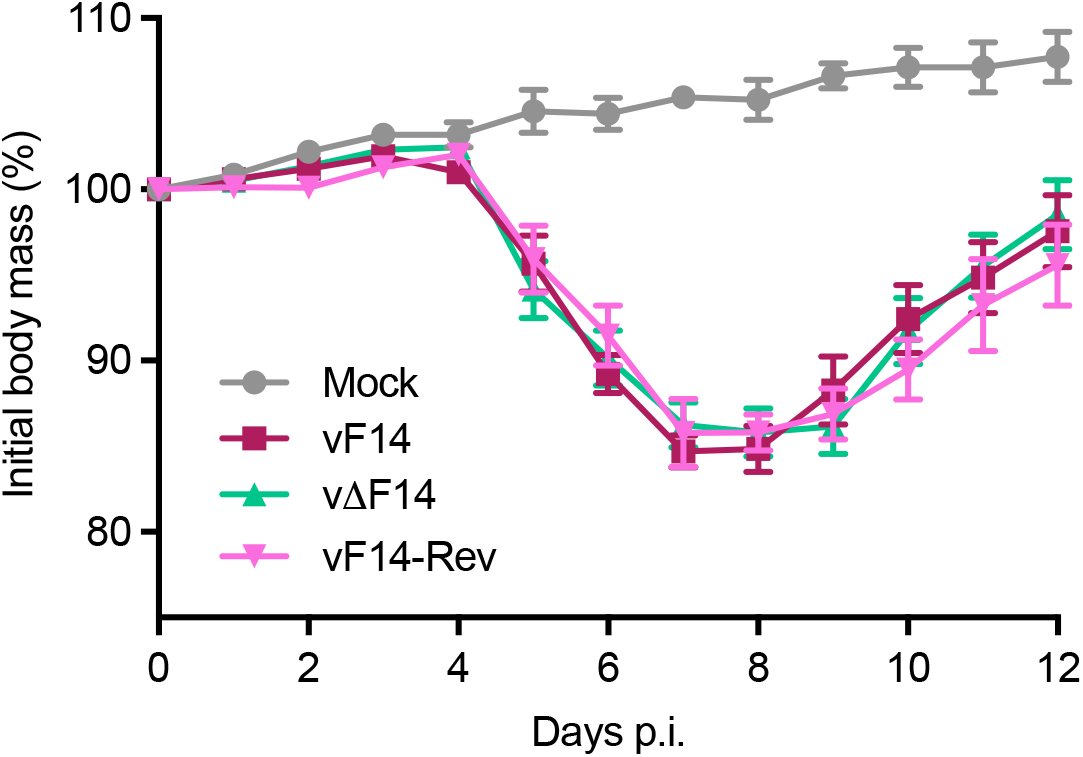
related to Figure 1: Virulence of VACV mutant lacking F14 in the intranasal mouse model of infection. BALB/c mice were infected intranasally with 5×10^3^ p.f.u. of the indicated VACV strains and their body mass was measured daily. Body mass is expressed as the percentage ± s.e.m. of the mean of the same group of mice on day 0 (*n* = 10 mice).

**Figure S3.**
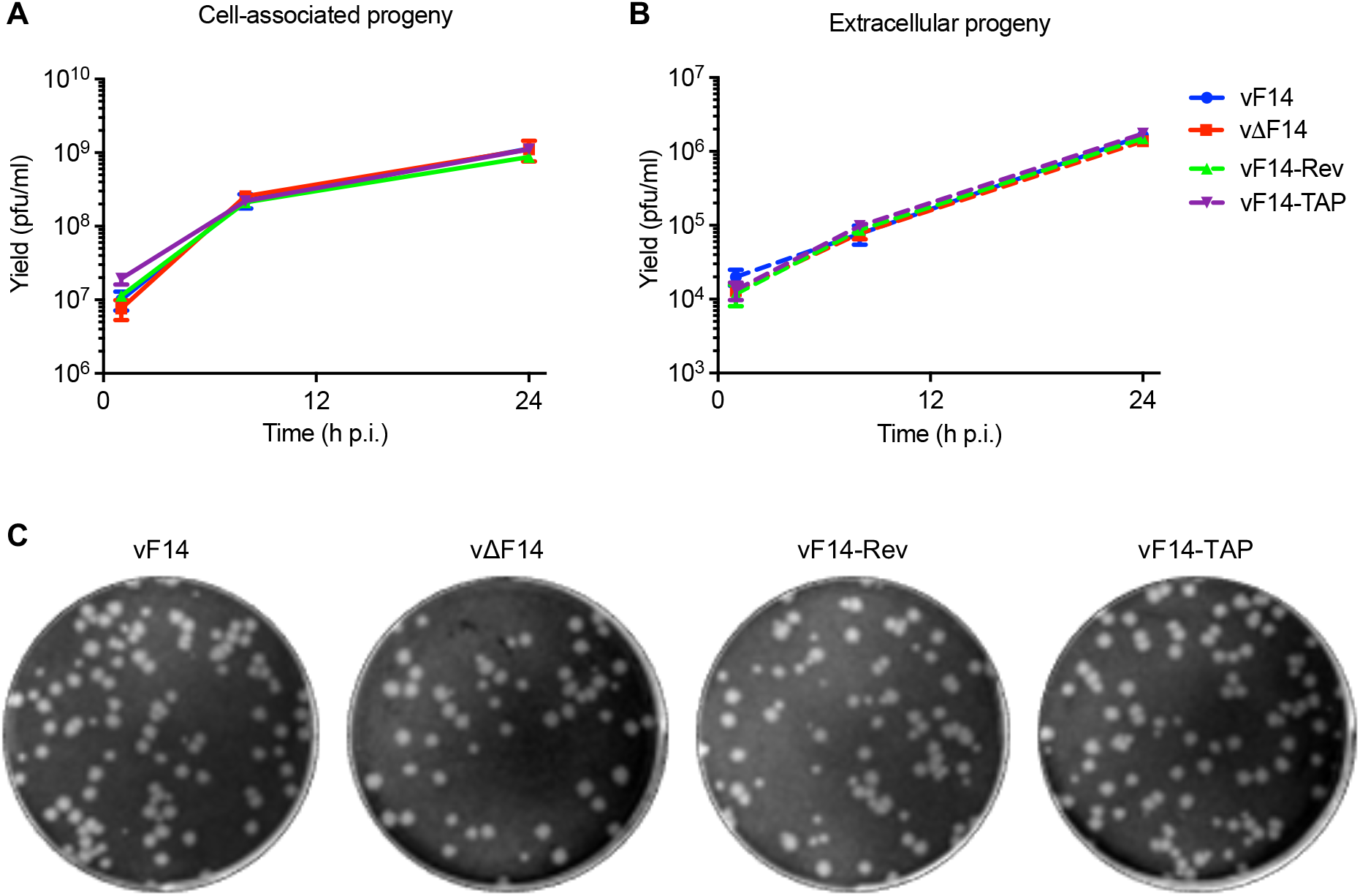
related to Figure 1: Replication and spread of VACV mutant lacking F14 in cell culture. (**A, B**) HeLa cells were infected with the indicated VACV strains (5 p.f.u./cell) and virus titres associated with the cells (**A**) and in the supernatants (**B**) were determined by plaque assay. Means ± s.d. (n = 2 per condition) are shown. (C) Plaque formation by the indicated VACV strains on BS-C-1 cells.

**Figure S4.**
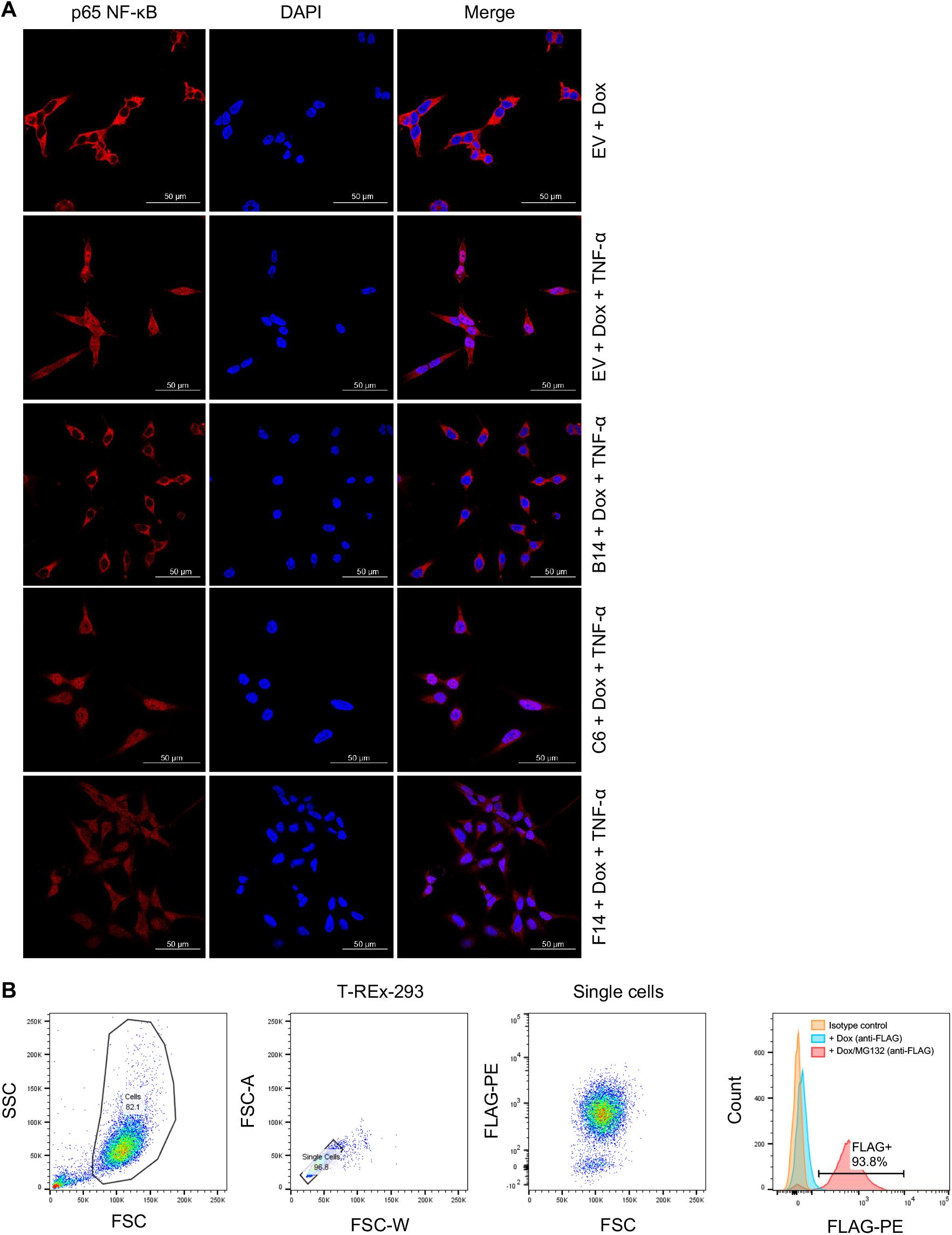
related to Figure 2: F14 does not inhibit the nuclear translocation of NF-κB subunit p65. **(A)** T-REx-293 cells inducibly expressing the empty vector (EV) or VACV proteins B14, C6, or F14, were induced with doxycycline and stimulated with TNF-α. Fixed and permeabilised cells were stained with anti-p65 antibody and DAPI, and analysed by confocal microscopy. Scale bars (50 μm) are shown in the bottom right of each microcraph. Representative micrographs of quantitative analysis shown in Figure 2B. **(B)** Flow cytometry analysis of T-REx-293-F14 induced with doxycycline in the absence and in the presence of the proteasome inhibitor MG132. F14 presence was detected by staining with an anti-FLAG antibody.

**Figure S5.**
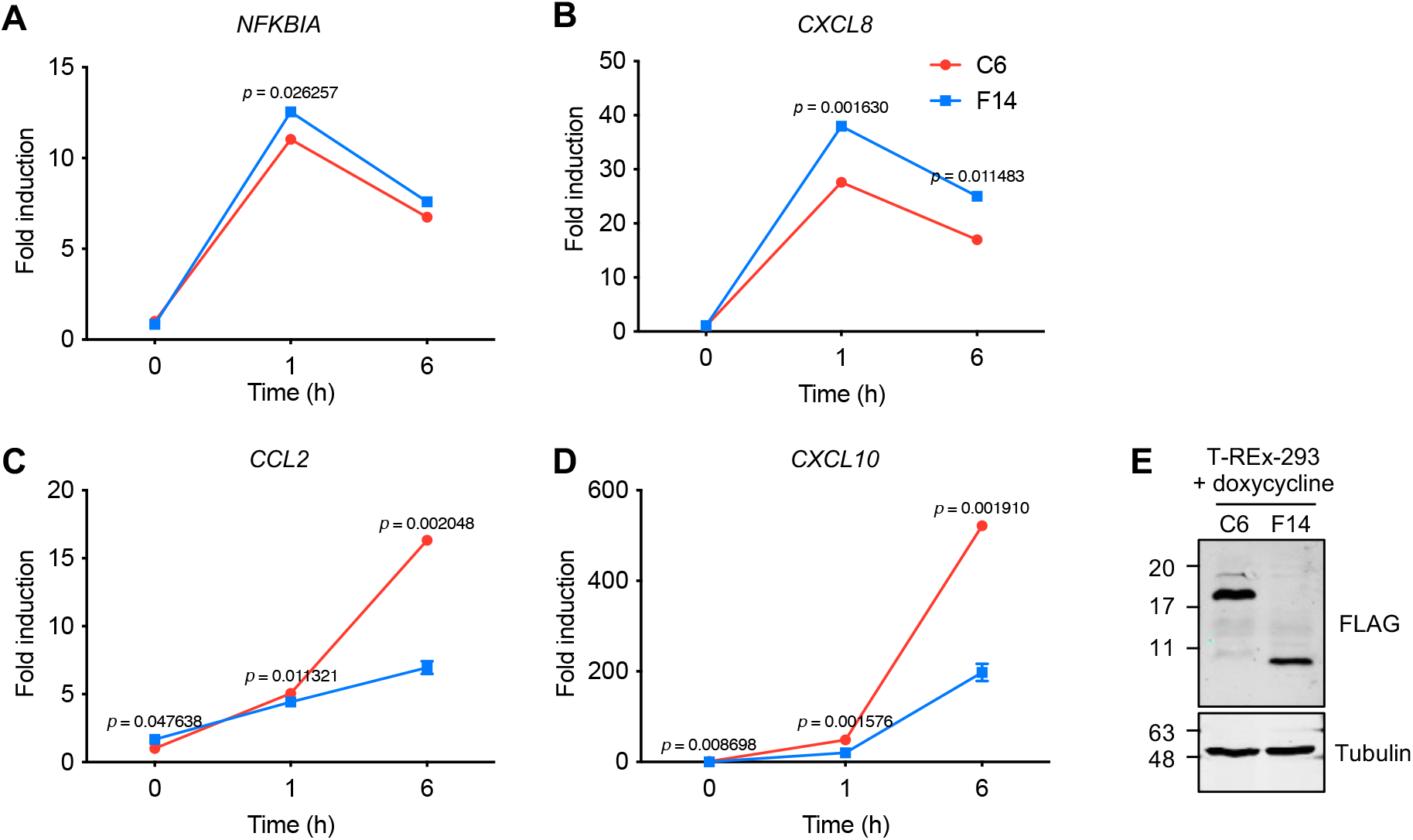
related to Figure 6: F14 suppresses expression of a subset of NF-κB- responsive genes. (**A-D**) RT-qPCR analysis of NF-κB-responsive gene expression in inducible T-REx-293 cells induced with doxycycline overnight to express VACV proteins F14 or C6, and stimulated with TNF-α. Means ± s.d. (*n* = 2 per condition) are shown. Statistical significance was determined by the Student’s *t*-test. (**E**) Immunoblotting of lysates of inducible T-REx-293 cell lines induced with doxycycline overnight. Protein molecular masses in kDa are shown on the left of the blots.

**Figure S6.**
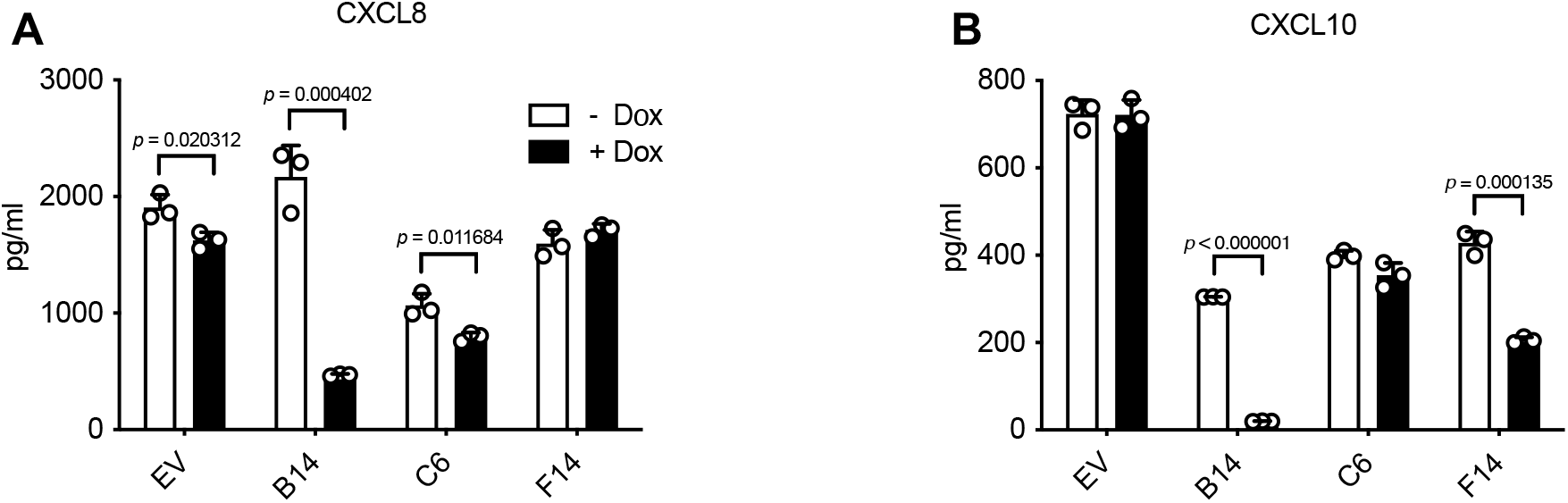
related to Figure 6: F14 suppresses expression of CXCL10, but not CXCL8, after stimulation with TNF-α. This shows data normalised for presentation in Figure 6C, F. ELISA of culture supernatants from T-REx-293 cells inducibly expressing the empty vector (EV) or VACV proteins B14, C6, or F14, induced with doxycycline and stimulated with TNF-α. Means + s.d. (*n* = 3 per condition) are shown. Statistical significance was determined by the Student’s *t*-test.

**Figure S7.**
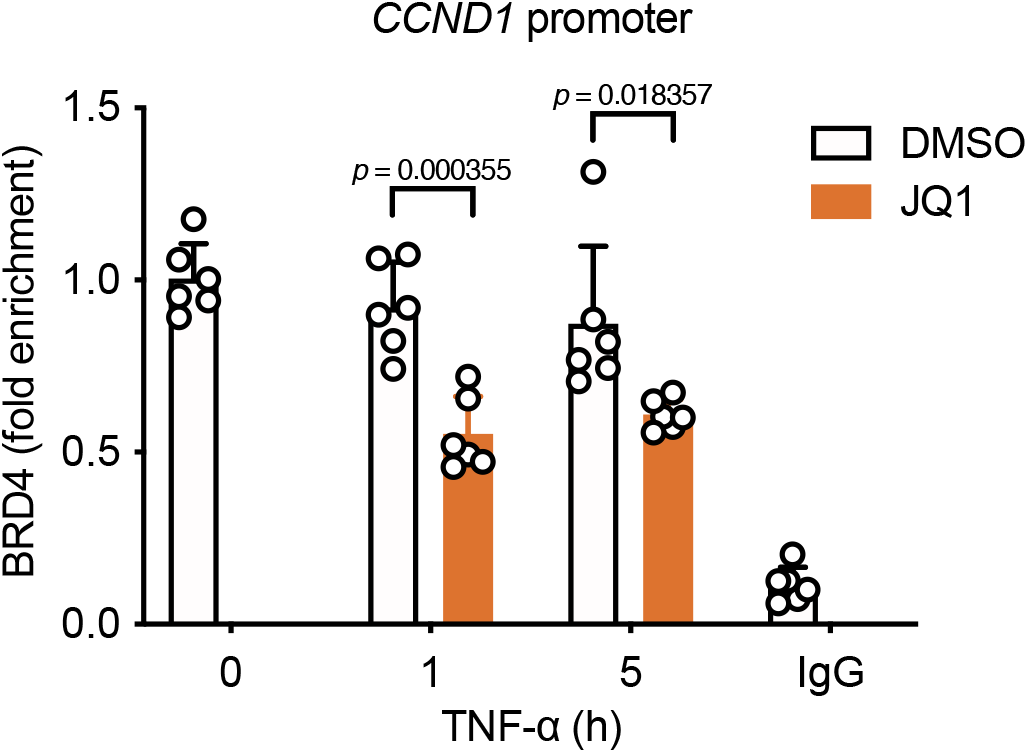
related to Figure 7: JQ1 reduces BRD4 occupancy on *CCND1* gene promoter. Chromatin immunoprecipitation (ChIP) with anti-BRD4 antibody or control IgG, and qPCR for the promoters of *CCND1* gene. T-REx-293 cells were treated with JQ1 and stimulated with TNF-α. Means + s.d. (*n* = 6 per condition from two independent experiments). Statistical significance was determined by the Student’s *t*-test.

**Table S1: NCBI GenBank accession numbers of poxvirus nucleotide sequences mentioned in this study.**

**Table S2: Oligonucleotide primers used in this study.** Primers are listed 5’ to 3’. Restriction sites used are highlighted in red and indicated in parentheses following oligonucleotide sequence. If present, sequences coding the tag epitopes are highlighted in bold, whilst the Kozak sequence are shown in italics. Plasmids marked with an asterisk (*) were constructed by site-directed mutagenesis, with the mutated codons underlined.

**Table S3: Antibodies used in this study.**

