## Supplemental Table 1 for "Viral mimicry of p65/RelA transactivation domain to inhibit NF-κB activation"

| **Virus species** | **Virus strain** | **NCBI accession number** |
|---|---|---|
| *Vaccinia virus* | Western Reserve | NC_006998.1 |
| *Vaccinia virus* | Copenhagen | M35027.1 |
| *Vaccinia virus* | Modified Virus Ankara, Acambis 3000 | AY603355.1 |
| *Vaccinia virus* | horsepox virus isolate MNR-76 | DQ792504.1 |
| *Monkeypox virus* | Zaire-96-I-16 | NC_003310.1 |
| *Cowpox virus* | Brighton Red | NC_003663.2 |
| *Variola virus* | India-1967 | NC_001611.1 |
| *Camelpox virus* | CMS | AY009089.1 |
| *Taterapox virus* | Dahomey 1968 | DQ437594.1 |
| *Ectromelia virus* | Moscow | AF012825.2 |
| *Raccoonpox virus* | Herman | NC_027213.1 |
| *Variola virus* | premodern variola virus, 10th century CE | LR800247.1 |
| *Variola virus* | premodern variola virus, 10th century CE | LR800244.1 |
| *Variola virus* | premodern variola virus, 10th century CE | LR800245.1 |
| *Variola virus* | premodern variola virus, 10th century CE | LR800246.1 |
| *Variola virus* | VD21, 17th century CE | KY358055.1 |

Table S1. NCBI GenBank accession numbers of poxvirus nucleotide sequences mentioned in this study.
