## Supplemental Table 1 for "Viral mimicry of p65/RelA transactivation domain to inhibit NF-κB activation"

| Target or Plasmid | Primers (5' to 3') | Reference |
|---|---|---|
| *CXCL8* (Fwd) | AGAAACCACCGGAAGGAACCATCT | Pallett et al. (2019) |
| *CXCL8* (Rev) | AGAGCTGCAGAAATCAGGAAGGCT | Pallett et al. (2019) |
| *NFKBIA* (Fwd) | CTCCGAGACTTTCGAGGAAAT | This study |
| *NFKBIA* (Rev) | GCCATTGTAGTTGGTAGCCTT | This study |
| *CCL2* (Fwd) | CAGCCAGATGCAATCAATGCC | Sumner et al. (2014) |
| *CCL2* (Rev) | TGGAATCCTGAACCCACTTCT | Sumner et al. (2014) |
| *CXCL10* (Fwd) | GTGGCATTCAAGGAGTACCTC | This study |
| *CXCL10* (Rev) | GCCTTCGATTCTGGATTCAGACA | This study |
| *GAPDH* (Fwd) | ACCCAGAAGACTGTGGATGG | Sumner et al. (2014) |
| *GAPDH* (Rev) | TTCTAGACGGCAGGTCAGGT | Sumner et al. (2014) |
| *NFKBIA* promoter (Fwd) | TCGCAGGGAGTTTCTCCGAT | This study |
| *NFKBIA* promoter (Rev) | TCAGGCTCGGGGAATTTCC | Tian et al. (2005) |
| *CXCL8* promoter (Fwd) | CATCAGTTGCAAATCGTGGA | Xu et al. (2012) |
| *CXCL8* promoter (Rev) | TGCACCCTCATCTTTTCATT | Xu et al. (2012) |
| *CCL2* promoter (Fwd) | CCCATTTGCTCATTTGGTCTCAGC | Harris et al. (2014) |
| *CCL2* promoter (Rev) | GCTGCTGTCTCTGCCTCTTATTGA | Harris et al. (2014) |
| *CXCL10* promoter (Fwd) | AGGAGCAGAGGGAAATTCCGTAAC | Harris et al. (2014) |
| *CXCL10* promoter (Rev) | AACGTGGGGCTAGTGTGCCA | Harris et al. (2014) |
| *CCND1* promoter (Fwd) | CTGAGCCGGGAAATCAACGA | This study |
| *CCND1* promoter (Rev) | ATAAAGAGGCTCGCCCACTC | This study |
| F14L sequencing primer (Fwd) | CGCAGGTACCGATGCAAATG | This study |
| F14L sequencing primer (Rev) | TGCTAGAATCAAGGAGGAGCA | This study |
| pcDNA4/TO-coF14-D62/63A-TAP* | CGAGAACTTCTCCGATATCGCCGCCGCCATCCTGGAAAGCCTGATCGAGC | This study |
| | GCTCGATCAGGCTTTCCAGGATGGCGGCGGCGATATCGGAGAAGTTCTCG | This study |
| pcDNA4/TO-coF14-D62/63K-TAP* | CGAGAACTTCTCCGATATCGCCAAGAAGATCCTGGAAAGCCTGATCGAGC | This study |
| | GCTCGATCAGGCTTTCCAGGATCTTCTTGGCGATATCGGAGAAGTTCTCG | This study |
| pcDNA4/TO-coF14-L65A-TAP* | CGATATCGCCGATGACATCGCCGAAAGCCTGATCGAGCAGG | This study |
| | CCTGCTCGATCAGGCTTTCGGCGATGTCATCGGCGATATCG | This study |
| pcDNA4/TO-coF14-L68A-TAP* | CGATGACATCCTGGAAAGCGCCATCGAGCAGGACGTGGCGG | This study |
| | CCGCCACGTCCTGCTCGATGGCGCTTTCCAGGATGTCATCG | This study |
| pcDNA4/TO-p65ΔTA1-TAP | AAAGGATCC*GCCACCATGG*ACGAACTGTTCCC (*BamHI*) | This study |
| | TTTGCGGCCGCGGCCCCCAGTGGAGCAGGAGCT (*NotI*) | This study |
| pcDNA4/TO-p65ΔTA1-F1451-73-TAP | AAAGGATCC*GCCACCATGG*ACGAACTGTTCCC (*BamHI*) | This study |
| | AAAGCGGCCGCCACGTCCTGCTCGATCAGGCTTTCCAGGATGTCATCGGCGATATCGGAGAAGTTCTCGTCGAAGTCCACGGCCCCCAGTGGAGCAGGAGC (*NotI*) | This study |
| pcDNA4/TO-coF14-FLAG | AAAGGATCC*GCCACCATG*AAGCACAGACTGTACAGCGA (*BamHI*) | This study |
| | AAAGCGGCCGCCACGTCCTGCTCGATCAGGC (*NotI*) | This study |
| pcDNA4/TO-coF6-FLAG | AAAGGATCC*GCCACCATG*AGCAAGATCCTGACCTTCGT (*BamHI*) | This study |

| | | |
|---|---|---|
| | AAAGCGGCCGCGTTGATGGTGAAGAAGCTCT (*NotI*) | This study |
| pcDNA4/TO-FLAG-coF7-WR | AAAGGATCC*GCCACCATGG*GCTCTTGCTGCGGCAGATT (*BamHI*) | This study |
| | AAATCTAGATTA**CTTGTCGTCGTCGTCCTTGTAGTC**CGCGGCCGCGAAGTACACGCTCTCGATTG (*XbaI* and *NotI*, respectively) | This study |
| pcDNA4/TO-coF7-Cop-FLAG | AAAGGATCC*GCCACCATGG*GCTTCTGCTGCGGCAGATT (*BamHI*) | This study |
| | AAAGCGGCCGCGAAGTACACGCTCTCGATTG (*NotI*) | This study |
| pcDNA4/TO-coB11-WR-FLAG | AAAGGATCCGCCACCATGGACACCGATGTGACCAACGT (*BamHI*) | This study |
| | AAAGCGGCCGCGATGCCCACGTCGTGGCAGA (*NotI*) | This study |
| pcDNA4/TO-coB11-Cop-FLAG | AAAGGATCCGCCACCATGGACACCGACACCGATACAGA (*BamHI*) | This study |
| | AAAGCGGCCGCGATGCCCACGTCGTGGCTGA (*NotI*) | This study |
| pcDNA4/TO-VP16-HA | TTTGGATCC*GCCACCATGG*ACCTCTTGG (*BamHI*) | This study |
| | TTTGCGGCCGCCCCACCGTACTCGTCAATTCC (*NotI*) | This study |
| pcDNA4/TO-VP16ΔTAD-HA | TTTGGATCC*GCCACCATGG*ACCTCTTGG (*BamHI*) | This study |
| | TTTGCGGCCGCCGTCGACAGTCTGCGCGTG (*NotI*) | This study |
| pcDNA4/TO-16E7-TAP | TTTGGATCC*GCCACCATG*CATGGAGATACACC (*BamHI*) | This study |
| | TTTGCGGCCGCTGGTTTCTGAGAACAGATGG (*NotI*) | This study |
| pcDNA4/TO-16E7-D75K-TAP* | TGCGTACAAAGCACACACGTAAAGATTCGTACTTTGGAAGACCTG | This study |
| | CAGGTCTTCCAAAGTACGAATCTTTACGTGTGTGCTTTGTACGCA | This study |
| pcDNA4/TO-16E7-L83R-TAP* | ATTCGTACTTTGGAAGACCTGAGAATGGGCACACTAGGAATTGTG | This study |
| | CACAATTCCTAGTGTGCCCATTCTCAGGTCTTCCAAAGTACGAAT | This study |
| pUC13-Ecogpt-EGFP-vΔF14 | AAACTGCAGATCAAAAATAAGCGCTCCCC (*PstI*) | This study |
| | AAAGCGGCCGCGAAGCTTAGGTTGTGATGTCGACTTTG (*NotI* and *HindIII*, respectively) | This study |
| | AAAAAGCTTCGCGGCCGCCATTGTAAATTTATAGGCGG (*HindIII* and *NotI*, respectively) | This study |
| | AAAGGATCCGGAAGCCTAAATTCG (*BamHI*) | This study |
| pUC13-Ecogpt-EGFP-vF14-Rev | AAACTGCAGATCAAAAATAAGCGCTCCCC (*PstI*) | This study |
| | AAAGGATCCGGAAGCCTAAATTCG (*BamHI*) | This study |
| pUC13-Ecogpt-EGFP-vF14-TAP | AAACTGCAGATCAAAAATAAGCGCTCCCC (*PstI*) | This study |
| | **GGCAGCGGAGGCGGAAGCTGGAGCCACCCCCAGTTCGAAAAGGGAGCCAGCGGCGAGGACTACAAGGATGACGATGACAAG**TAAGTTTTTATGTTAACTAA | This study |
| | **TCCCTTTTCGAACTGGGGGTGGCTCCAGCTTCCGCCTCCGCTGCCTCCTCCCTTTTCAAACTGAGGATGAGACCA**CGCGGCCGCTACATCCTGTTCTATCA (*NotI*) | This study |
| | AAAGGATCCGGAAGCCTAAATTCG (*BamHI*) | This study |
| pUC13-Ecogpt-EGFP-vv811ΔA49ΔF14 | AAACTGCAGATCAAAAATAAGCGCTCCCC (*PstI*) | This study |
| | AAAGCGGCCGCGAAGCTTAGGTTGTGATGTCGACTTTG (*NotI* and *HindIII*, respectively) | This study |

| | | |
|---|---|---|
| | AAAAAGCTTCGCGGCCGCCATTGTAAATTTATAGGCGG (*HindIII* and *NotI*, respectively) | This study |
| | AAAGGATCCGGAAGCCTAAATTCG (*BamHI*) | This study |

Table S2. Oligonucleotide primers used in this study. Primers are listed 5’ to 3’. Restriction sites used are highlighted in red and indicated in parentheses following oligonucleotide sequence. If present, sequences coding the tag epitopes are highlighted in bold, whilst the Kozak sequence are shown in italics. Plasmids marked with an asterisk (*) were constructed by site-directed mutagenesis, with the mutated codons underlined.
