## Supplemental Table 1 for "Viral mimicry of p65/RelA transactivation domain to inhibit NF-κB activation"

| ANTIBODY | DILUTION (IB/IF/FC) OR AMOUNT (ChIP)* | BLOCKING AGENT (IB/IF/FC)* | SOURCE | IDENTIFIER |
|---|---|---|---|---|
| Mouse monoclonal anti-FLAG (clone M2) | 1:1,000 (IB); 1:1,000 (IF); 1:650 (FC) | 5% non-fat milk (IB); 10% FBS* (IF); 0.1% BSA (FC) | Sigma-Aldrich | Cat# F3165; RRID:AB_259529 |
| Rabbit polyclonal anti-FLAG | 1:5,000 (IB); 1:2,000 (IF) | 5% non-fat milk (IB); 10% FBS* (IF) | Sigma-Aldrich | Cat# F7425; RRID:AB_439687 |
| Rat monoclonal anti-α-Tubulin (clone YL1/2) | 1:10,000 (IB) | 5% non-fat milk (IB) | Serotec | Cat# MCA77G; RRID:AB_325003 |
| Rabbit polyclonal anti-C6 | 1:1,000 (IB) | 5% non-fat milk (IB) | Laboratory of Geoffrey Smith; Unterholzner et al., 2011 | N/A |
| Mouse monoclonal anti-D8 (clone AB1.1) | 1:1,000 (IB) | 5% non-fat milk (IB) | Laboratory of Geoffrey Smith; Parkinson and Smith, 1994 | N/A |
| Rabbit polyclonal anti-A49 | 1:1,000 (IB) | 5% non-fat milk (IB) | Laboratory of Geoffrey Smith; Mansur et al., 2013 | N/A |
| Rabbit polyclonal anti-VACV | 1:1,000 (IB) | 5% non-fat milk (IB) | Laboratory of Geoffrey Smith; unpublished | N/A |
| Mouse monoclonal anti-IκBα (L35A5) | 1:1,000 (IB) | 5% non-fat milk (IB) | Cell Signaling Technology | Cat# 4814; RRID:AB_390781 |
| Rabbit polyclonal anti-phospho-p65 (Ser536) | 1:1,000 (IB) | 3% BSA* | Cell Signaling Technology | Cat# 3031; RRID:AB_330559 |
| Rabbit monoclonal anti-p65 (D14E12) | 1:1,000 (IB); 1:400 (IF) | 3% BSA* (IB); 10% FBS* (IF) | Cell Signaling Technology | Cat# 8242; RRID:AB_10859369 |
| Mouse monoclonal anti-p65 (F-6) | 1:1,000 (IB); 1:100 (IF) | 5% non-fat milk (IB); 10% FBS* (IF) | Santa Cruz Biotechnology | Cat# sc-8008; RRID:AB_628017 |
| Rabbit polyclonal anti-HA | 1:1,000 (IB) | 5% non-fat milk (IB) | Sigma-Aldrich | Cat# H6908; RRID:AB_260070 |
| Rabbit polyclonal anti-CBP (A-22) | 1:500 (IB) | 5% non-fat milk (IB) | Santa Cruz Biotechnology | Cat# sc-369; RRID:AB_631006 |
| Rabbit monoclonal anti-acetyl-p65 (Lys310) (D2S3J) | 1:1,000 (IB) | 3% BSA* (IB) | Cell Signaling Technology | Cat# 12629; RRID:AB_2722509 |
| Rabbit monoclonal anti-BRD4 (E2A7X) | 8.0 µg (ChIP) | NA | Cell Signaling Technology | Cat# 13440; RRID:AB_2687578 |
| Rabbit polyclonal anti-GFP | 8.0 µg (ChIP) | NA | Abcam | Cat# ab290; RRID:AB_303395 |
| Mouse monoclonal anti-GAPDH (clone 71.1) | 1:1,000 (IB) | 5% non-fat milk (IB) | Sigma-Aldrich | Cat# G8795; RRID:AB_1078991 |
| IRDye 680RD-conjugated goat anti-rabbit IgG | 1:10,000 (IB) | 5% non-fat milk (IB) | LI-COR | Cat# 926-68071; RRID:AB_10956166 |
| IRDye 680LT-conjugated goat anti-mouse IgG | 1:10,000 (IB) | 5% non-fat milk (IB) | LI-COR | Cat# 926-68020; RRID:AB_10706161 |
| IRDye 800CW-conjugated goat anti-rabbit IgG | 1:10,000 (IB) | 5% non-fat milk (IB) | LI-COR | Cat# 926-32211; RRID:AB_621843 |
| IRDye 800CW-conjugated goat anti-mouse IgG | 1:10,000 (IB) | 5% non-fat milk (IB) | LI-COR | Cat# 926-32210; RRID:AB_621842 |
| IRDye 680LT-conjugated goat anti-rat IgG | 1:10,000 (IB) | 5% non-fat milk (IB) | LI-COR | Cat# 926-68029; RRID:AB_10715073 |
| Donkey anti-mouse IgG (H+L) secondary antibody, Alexa Fluor 546 | 1:1,000 (IF) | 10% FBS* (IF) | Molecular Probes | Cat# A10036; RRID:AB_2534012 |
| Goat anti-rabbit IgG (H+L) secondary antibody, Alexa Fluor 488 | 1:1,000 (IF) | 10% FBS* (IF) | Molecular Probes | Cat# A11008; RRID:AB_143165 |
| PE goat anti-mouse IgG (clone Poly4053) | 1:100 (FC) | 0.1% BSA (FC) | BioLegend | Cat#405307; RRID:AB_315010 |

*IB, immunoblotting; IF, immunofluorescence; ChIP, chromatin immunoprecipitation; FC, flow cytometry; FBS, foetal bovine serum; BSA, bovine serum albumin.

Table S3. Antibodies used in this study.
